# Naturally segregating genetic variants contribute to thermal tolerance in a *D. melanogaste*r model system

**DOI:** 10.1101/2023.07.06.547110

**Authors:** Patricka A. Williams-Simon, Camille Oster, Jordyn A. Moaton, Ronel Ghidey, Enoch Ng’oma, Kevin M. Middleton, Troy Zars, Elizabeth G. King

## Abstract

Thermal tolerance is a fundamental physiological complex trait for survival in many species. For example, everyday tasks such as foraging, finding a mate, and avoiding predation, are highly dependent on how well an organism can tolerate extreme temperatures. Understanding the general architecture of the natural variants of the genes that control this trait is of high importance if we want to better comprehend how this trait evolves in natural populations. Here, we take a multipronged approach to further dissect the genetic architecture that controls thermal tolerance in natural populations using the Drosophila Synthetic Population Resource (DSPR) as a model system. First, we used quantitative genetics and Quantitative Trait Loci (QTL) mapping to identify major effect regions within the genome that influences thermal tolerance, then integrated RNA-sequencing to identify differences in gene expression, and lastly, we used the RNAi system to 1) alter tissue-specific gene expression and 2) functionally validate our findings. This powerful integration of approaches not only allows for the identification of the genetic basis of thermal tolerance but also the physiology of thermal tolerance in a natural population, which ultimately elucidates thermal tolerance through a fitness-associated lens.

## Introduction

Thermal tolerance is a complex physiological and behavioral trait that determines the survivability of many individuals and, ultimately populations. Temperature regulates and plays a critical role in a wide range of processes from cellular functions – action potentials and enzymatic reactions (Schulte *et al*. 2011; Yu *et al*. 2012; Dowd *et al*. 2015) – to more systematic evolutionary processes like migration and population growth (Frazier *et al*. 2006; Sunday *et al*. 2012) and ultimately fitness (Angilletta and Angilletta 2009). Indeed, any change in optimum temperature can cause multiple negative downstream effects on cellular and population levels (Chen *et al*. 2011; Huey *et al*. 2012; Sunday *et al*. 2012; Urban 2015). This is especially concerning as climate change and increased frequency of extreme temperature events continue to threaten our planet (McCain and Colwell 2011; Letcher 2015) and force the redistribution and/or adaptation of many organisms and reviewed in (Walther *et al*. 2002; Parmesan and Yohe 2003; Thomas 2010; Bozinovic *et al*. 2011; Sunday *et al*. 2012). Climate change is a major ecological and evolutionary problem (Huey *et al*. 2009; Harley and Paine 2009; Allen *et al*. 2010; McKechnie and Wolf 2010; Kjærsgaard *et al*. 2010; Denny and Dowd 2012; Kingsolver and Buckley 2015; Buckley and Huey 2016; Seuront *et al*. 2019) and continues to be one of the leading causes of species extinction (Thomas *et al*. 2004; Warren *et al*. 2013; Román-Palacios and Wiens 2020) reviewed in (Pimm 2009; Cahill *et al*. 2013). Therefore, having a comprehensive understanding of the genetic architecture that underlies the genomic regions that influence thermal tolerance is fundamental to elucidating complex fitness-associated processes, such as evolutionary rescue. This understanding is expected to provide insights into the individuals, and ultimately, populations that will be the ‘winners and losers’ (Somero 2010, 2011).

Many empirical studies – both in model and nonmodel systems – demonstrate the importance of understanding the mechanisms contributing to thermal tolerance, including in coral (Dixon *et al*. 2015), fish (Schulte *et al*. 2011; Anttila *et al*. 2013; Grinder *et al*. 2020), fruit flies (Overgaard *et al*. 2008; Bozinovic *et al*. 2016), birds (Nord and Nilsson 2011; DuRant *et al*. 2012, 2013), frogs (Pintanel *et al*. 2019; Díaz-Ricaurte *et al*. 2021), and lizards (Angilletta *et al*. 2002a). The ability to tolerate extreme temperatures is essential for ectotherms (Huey and Berrigan 2001; Angilletta *et al*. 2002b; Frazier *et al*. 2006; Hoffmann 2010; Sunday *et al*. 2011; Gunderson and Stillman 2015) because their body temperature tracks the ambient temperature to a greater degree than endotherms. *Drosophila* species have been used extensively as a research model because of their worldwide distribution and vastly different thermal environments and preferences (MacLean *et al*. 2019a). *D. melanogaster* in particular, has been used for decades to study these traits in laboratory and field settings, (James *et al*. 1997; Hoffmann *et al*. 2003; Kristensen *et al*. 2008; Bergland *et al*. 2014; Ørsted *et al*. 2019; Machado *et al*. 2021; Rudman *et al*. 2022). For example, within a natural environment, Machado et al., (2021), demonstrated the effects of seasonal temperature shifts in allele frequencies of fruit flies, suggesting that short-term temperature changes can lead to rapid changes in allele frequencies. While these studies have been imperative in elucidating how ectotherms adapt to temperature changes and how the genome responds to such changes, there is still a lack of complete knowledge about the specific genetic mechanisms underpinning thermal tolerance.

Identifying the causative mechanisms at play in highly complex, multifaceted traits like thermal tolerance remains very challenging. Genetic mapping approaches, designed to identify loci associated with the naturally occurring variation in a trait continue to improve and have led to the discovery of many associations for thermal tolerance in fruit flies (Morgan and Mackay 2006; Vermeulen *et al*. 2008a; b; Rand *et al*. 2010; Rolandi *et al*. 2018) and other systems including coral, fish, pigs, and cattle (Perry *et al*. 2001; Everett and Seeb 2014; Dixon *et al*. 2015; Matz *et al*. 2018; Kim *et al*. 2018; Cheruiyot *et al*. 2021). However, mapping approaches lack the resolution to identify single genes on their own (Mackay 2001). More recently researchers have been pairing mapping approaches with transcriptomics to identify candidate genes that influence complex traits (Cubillos *et al*. 2017; Highfill *et al*. 2017; Williams-Simon *et al*. 2019; Wen *et al*. 2019; Ng’oma *et al*. 2020; Lecheta *et al*. 2020), which was first introduced by Wayne and McIntyre (2002). While it is critically important to validate candidate genes that have been identified to be associated with complex traits, it has been a daunting task, and therefore few genetic studies of complex traits have definitively identified causal variants (Ding *et al*. 2016; Highfill *et al*. 2017; Gaspar *et al*. 2020). Approaches integrating several methods including multiparent mapping for high power and resolution, transcriptomics, and functional genomics provide a promising approach that moves beyond simply mapping loci toward identifying the key genes involved in complex traits.

Here, we leverage the combined strengths of QTL mapping, transcriptomics, and functional genetics to elucidate the genetic architecture of thermal tolerance using a large set of recombinant inbred lines (RIL) from the *Drosophila* Synthetic Population Resource (DSPR) multi-parental population (King *et al*. 2012a, MPP; b). We measured over 700 RILs for thermal tolerance and identified multiple QTLs, including one large effect QTL. We then performed RNA-seq to identify several candidate genes that are located within our QTLs and performed validation studies using classical genetics. Through this multifaceted approach, we identified novel candidate genes associated with the critically important trait, thermal tolerance.

## Methods

### DSPR and Fly Husbandry

We used the DSPR, which is a multi-parental mapping population for the *D. melanogaster* system, (King *et al*. 2012a; b; Long *et al*. 2014); http://FlyRILs.org). The DSPR consists of approximately 1,600 8-way Recombinant Inbred Lines (RILs), with two populations (pA and pB), which consist of approximately 800 RILs each. Each population was created by crossing a set of 8 founder lines, with seven unique founders to each population and one that is shared – AB8. Each RIL is a mosaic of the 15 founders (parental) lines, which have been fully re-sequenced, and the RILs have been genotyped at over 10,000 SNPs. This structure makes genomic analysis of this population more feasible, because of the ability to be able to track entire haplotypes (King *et al*. 2012a). To generate the RILs, the founders were mixed *en masse* for 50 generations to create a synthetic population, with two major goals - first to generate high genetic diversity, then for high mapping resolution. The population was then inbred for another 25 generations to create stable recombinant inbred lines (King *et al*. 2012b). Within each of the pA and pB populations, two separate synthetic populations were maintained (e.g., pA1 and pA2) for the crossing phase before the creation of the RILs. We phenotyped 741 RILs in population A, scoring thermal tolerance for each line (Fig. 1A).

**Figure 1:**
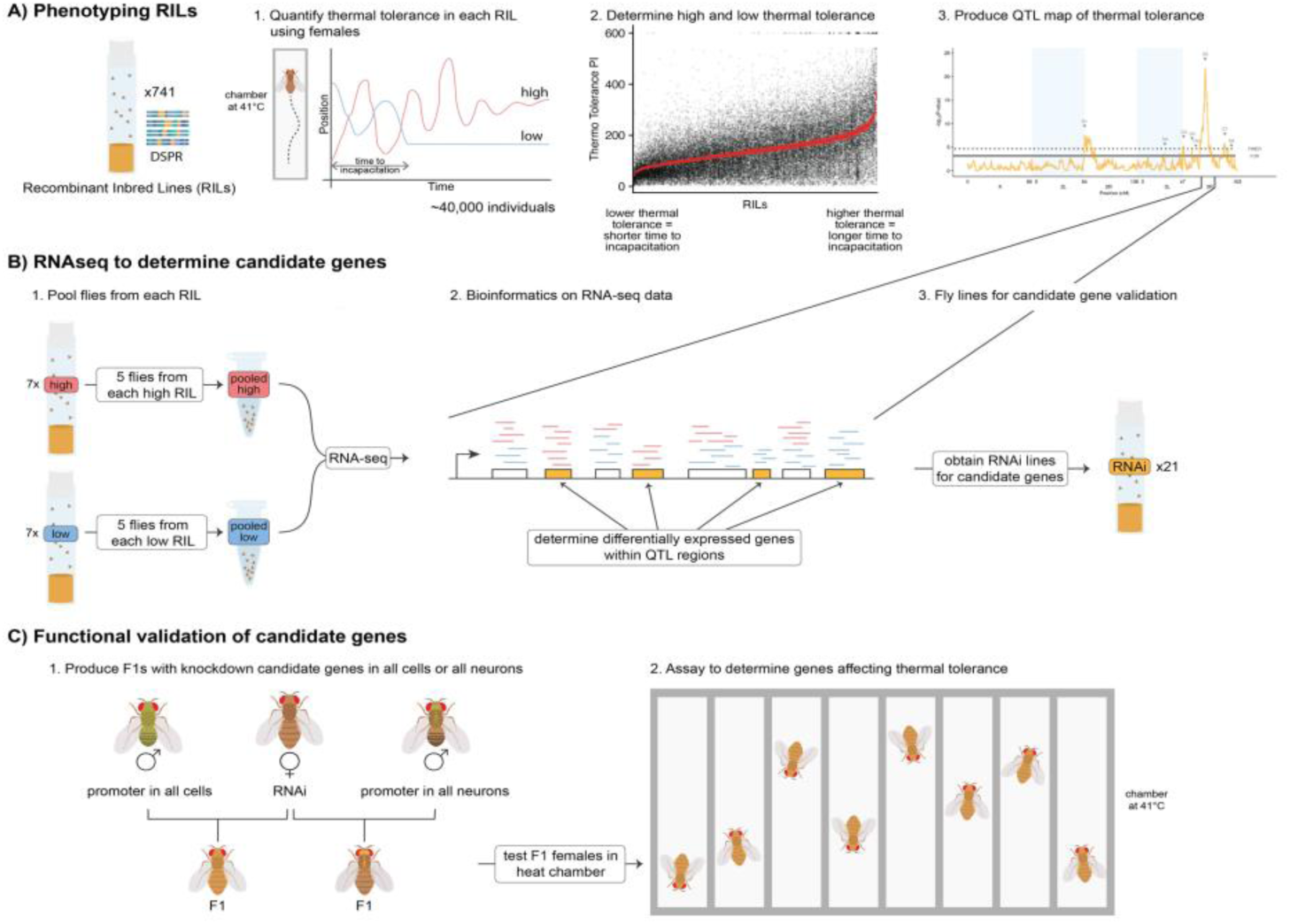
Schematic of the experimental design including A) phenotyping and QTL mapping using the DSPR RILs, B) RNAseq and differential expression analysis, and C) using RNAi lines for functional validation of candidate genes within the QTL interval.

We raised stock RILs at 18°C and 60% relative humidity on a standard fruit fly diet (Table S1) with live yeast added to the top of the food after it cooled. Two weeks before phenotyping for thermal tolerance, approximately 10 female and 6 male flies were placed into a flask (Fisher Scientific, Cat. No.:AS-355) of food and maintained at 25°C for one week to allow for mating and egg laying. After the first week (7 days, post-oviposition), we removed all the adults (parental generation) from the flasks to ensure that there was no mixing of generations. After two weeks, (14 days post-oviposition) we anesthetized adult flies on ice for no more than 10 minutes and selected 60-80 female flies for the thermal tolerance assay. We then divided flies equally between two separate vials of food and allowed them to recover for at least 24 hours before the phenotyping assay.

### Measuring Thermal Tolerance in the DSPR

We measured thermal tolerance using a thermally-controlled and highly sensitive apparatus known as the “heat box” (Wustmann *et al*. 1996). Within the heat box, there are 16 individual chambers, which allows for concurrent testing of flies. In each chamber, temperatures are set at 24°C for the first 30 secs for acclimation, then immediately shift to 41°C for 9.5 minutes for a total of 10 minutes (Wustmann *et al*. 1996; Zars 2001). We assayed the RILs in a randomized order, limiting the number of flies assayed on a single day from a single RIL up to 27 individuals total (two groups in the heat box), to ensure each RIL was assayed over multiple days, however, we ran several RILs on any given day. For each RIL, we assayed a minimum of 30 individuals. We also measured the founder lines for thermal tolerance, following the same experimental design as we did for the RILs.

We define thermal tolerance as the amount of time it takes before a fly becomes incapacitated for at least 60 seconds. We used a custom R script to automate scoring incapacitation based on the criteria defined above (i.e., no activity for 60 seconds). The heat box contains an infrared system for monitoring each fly’s position within the chamber. We used a sliding window to assess the variance in position in 60 second intervals and scored a fly as incapacitated when this variance was below a threshold value indicating no substantial movement within that interval. This approach has the advantage of applying purely objective, defined criteria to scoring incapacitation. However, it is important to confirm the program produced the correct results. We validated our script by comparing over 5000 visually hand-scored incapacitation scores to our automated-scored values (Fig. S1). These scores are highly correlated (r = 0.88), and manual inspection of discrepancies between the hand-scored and automated-scored values revealed the automated values to be more accurate and consistent for our defined criteria for incapacitation.

### QTL Mapping and Heritability

We performed a genome scan on 741 RILs using a custom R script as described in Williams-Simon et al. (2019). We used the average incapacitation times from each RIL as phenotype and applied the Haley-Knott regression (Haley and Knott 1992). This statistical approach regresses each phenotypic average on the eight founder haplotype probabilities (Broman and Sen 2009; King *et al*. 2012a) by fitting the following model:

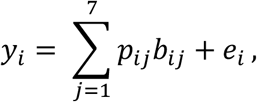

where *y*_*i*_ is the phenotype (i.e., thermal tolerance) of the ith RIL, *p*_*ij*_ is the probability the *it*ℎ RIL has the *jt*ℎ haplotype at the locus, *b*_*ij*_ is the vector of effects for the *jt*ℎ haplotype, and *e*_*i*_ is the vector of residuals (Broman and Sen 2009). Prior to performing this genome scan we transformed the raw phenotypic data, using a square root transformation, which improved the normality distribution of the data to meet the assumptions of our statistical test. The DSPR RILs contain two subpopulations within each set of RILs, stemming from the synthetic populations being maintained *en masse* in two sets before the creation of the RILs (King *et al*. 2012a). Including subpopulations (pA1 and pA2) in the model did not affect the mapping results (Fig. S2A). We also performed a second genome scan after correcting for our most significant QTL (Q5) and did not obtain any additional peaks (Fig. S2B).

To identify statistically significant QTLs, we used permutations to calculate both the false discovery rate (FDR) and the family-wise error rate (FWER). First, we permuted thermal tolerance along with place learning and memory phenotypes (Williams-Simon *et al*. 2019), because we assayed all three of these phenotypes on the same individual fly during the experiment. Second, we computed the number of false positive QTLs at a range of different significance thresholds. We removed any QTL that were within 2 cM of a more significant QTL to identify distinct QTLs. Then, we calculated the empirical false discovery rate (FDR, expected false positives/total positives) for each threshold and determined the threshold corresponding to a 5% FDR. We also calculated the threshold corresponding to a 5% family-wise error rate (FWER, the probability of one or more false positives experiment-wide) by determining the lowest P-value for each set of genome scans from each permutation and calculated the 95% quantile of the resulting set of P-values (Churchill and Doerge 1994). For each identified peak, we calculated the estimated effects of each haplotype at each QTL, the percent variance explained (PVE) by the QTL and the Bayesian credible interval (BCI), following the methods described previously for the DSPR (King *et al*. 2012a). If the BCI of two significant QTLs did not overlap, we considered these distinct QTLs.

We calculated broad sense heritability (H^2^ = V_G_/V_p_) of thermal tolerance by estimating the genetic and phenotypic variance and covariance from a linear mixed model *lme* and *VarCorr* functions in the *nlme* package (Pinheiro and Bates 2000; Pinheiro *et al*. 2023) followed by a jackknife, which allowed us to obtain standard errors of our estimate (Roff and Preziosi 1994; Roff 2008). The jackknife removes each observation once, the model is then fit, and the quantitative genetic parameter (i.e., heritability) is estimated and a pseudo value is calculated. Prior to fitting the model, we quantile-transformed the raw data to ensure normality. Thus, for our dataset of 741 RILs, each RIL is deleted, one at a time, to produce 741 pseudo values. Because the *lme* function uses restricted maximum likelihood, it is possible for the model to not converge. This occurred for 72 cases, thus, these pseudo values were not included in the calculation of our estimates. We also performed a model comparison using a likelihood ratio test to determine the significance of RIL (i.e., genotype). We compared the model fitting only the intercept to the model including RIL, fitting both via maximum likelihood such that model comparisons would be valid.

### Structural Variants Analysis

*Structural Variants (SVs)*, for example, insertions, deletions, and transposable elements contribute to the phenotypic variation in complex traits (Frazer *et al*. 2009; Eichler *et al*. 2010; Chakraborty *et al*. 2018). Chakraborty et al. (2019) used the DSPR to identify genome-wide SVs in both populations A and B, which we utilized to determine if there were any SVs located within the BCI of our most significant QTL (Q5). To ensure that we captured all the SVs located in the QTL region, we included any genes that had any portion within the BCI, even if the entire gene did not fall within the QTL BCI.

### RNA-sequencing

We performed RNA-seq on female heads to determine differentially expressed genes between the top and bottom cohorts of thermal tolerance (Fig. 1B). It is important to note that in our design we only used heads which allowed us to only focus on gene expression within a specific tissue type. It is suggested in the literature that chill coma recovery (thermal tolerance) in insects is controlled by different tissue types (Andersen and Overgaard 2019). First, we selected 14 RILs from the high (n = 7) and low (n = 7) 5% cohorts after phenotyping the first 140 RILs and followed the same set-up protocol (see phenotyping section above). In total, we had 6 pooled samples consisting of 5 individual female heads from each of the 7 lines in the cohort. Thus, for each of the 6 samples, we pooled a total of 35 female heads in a single sample, with 5 heads from each RIL contributing to each sample. Adult female flies 14 days post-oviposition were flash-frozen in liquid nitrogen and stored at -80°C, until further processing. In addition, we also took a different set of individuals from the same RILs and cycled them through an ‘incubation assay’ for 1 min intervals at 41°C and 24°C (room temp) for a total of 6 mins., and left control samples on the benchtop at a constant 24°C also for 6 minutes. Then, all the vials – from both experimental and control groups – were left on the benchtop at room temp for 3 hrs, to ensure that the gene expression changes had time to occur but also that there was minimal time to allow for gene expression turnover. Immediately after the 3 hrs, flies were flash-frozen in liquid nitrogen and stored at -80°C. For both the heat box and incubator assays, only female flies ranging from 15-22 days old post-oviposition were assayed. For the incubator assay, all RILs were tested between 8 am and 10 am, to help reduce possible confounding effects that would affect gene expression. The samples were then freeze-dried overnight to prevent the degradation of RNA, using a lyophilizer (*Labconco, Cat No.: 77550-00*). Second, the samples were shipped to Rapid Genomics (Gainesville, FL) for transcriptome processing. The total RNA was extracted with Dynabeads mRNA direct kit from life technologies, mRNA was then fragmented and converted into double-stranded cDNA, followed by the standard proprietary library prep for one lane of Illumina HiSeq 3000 instrument to generate paired-end (PE) reads. The first 16 samples were PE 150 bp, while the remaining samples were PE 100 bp. Third, We analyzed the RNA-seq data using a bioinformatic pipeline, as described previously in Williams-Simon et al. (2019). Briefly, we trimmed the samples using the software *cutadapt (Martin 2011)*, then ran a quality control test using *fastqc (Andrews and Others 2010),* which did not show any significant issues with the reads. We then used parts of the *Tuxedo* pipeline *(Pertea et al. 2016)* to align the reads against the *D. melanogaster* reference transcriptome (Ensembl Release Version: 84) using *HISAT2 (Kim et al. 2015)* and *StringTie (Pertea et al. 2015, 2016)* assembled and quantified transcripts. To calculate the read count for each gene we used a *prepDE* python script provided for this purpose (http://ccb.jhu.edu/software/stringtie/index.shtml?t=manual#deseq). The resulting dataset provided the input needed to use the *DESeq2* package (Love *et al*. 2014) to identify differentially expressed genes. We filtered out any zero count genes and performed surrogate variables analysis (SVA) on the normalized count data (Leek *et al*. 2012, 2019) to estimate any unknown batch effects. We identified 2 significant surrogate variables, which we included as terms in the models we fit along with our known batch effect (i.e., sampling day) in all following bioinformatics analyses. We then used the DESeq2 package (Love *et al*. 2014) to test for overall treatment effects and performed contrasts between pool (high and low cohorts), condition (temperature: 41°C or room temperature), and the interaction between pool and condition, identifying significantly differentially expressed genes for each. To estimate the differences in expression in each pool and condition, we used the function *lfcShrink*, which performs shrinkage on log_2_(Fold Changes). All log_2_(Fold Changes) reported here are the shrinkage estimated values using the “*normal*” estimator. We then used the contrasts to identify candidate genes that were differentially expressed between the different groups localized within the BCIs of the mapped QTLs (Q1-Q7).

### RNAi Lines and Crossing Design

Because the Q5 QTL indicated a very strong association with thermal tolerance and the BCI was fairly narrow, we further investigated the genes within this QTL using RNAi lines. We sourced the UAS-RNAi transgenic fly lines from the Transgenic RNAi Project (TRiP) from the Bloomington Drosophila Stock Center (BDSC; Perkins *et al*. 2015)). BDSC reported that the TRiP RNAi lines might have a low dose of *Wolbachia pipientis*, therefore we treated the flies with tetracycline (Sigma-Aldrich, Cat No.:T4062-5G, Batch: SLCF9393). We followed a tetracycline treatment protocol from Hoffman et al., (Hoffmann *et al*. 1986). We incorporated 0.3 mg/mL final concentration of tetracycline into an agar base fly food recipe and allowed the flies to remain on that food for two generations flipping them onto new agar food after 2 weeks. One of the lines did not survive during the tetracycline treatment (BDSC stock No.: 63713). After the treatment, we switched the population back to our ‘normal’ fly food recipe (Table S1) and allowed the lines to re-acclimate to the food for 5 generations before beginning experiments. The TRiP collection did not have RNAi lines for all 21 genes in the Q5 interval. However, we obtained RNAi lines for 14 genes from the BDSC, some of which had more than one available RNAi line (Table S2). We checked each UAS line for reports of off-target effects and did not identify any reported off-target effects (https://fgr.hms.harvard.edu/up-torr). In total we screened 20 UAS-RNAi lines using two different promoters: 1) ubiquitously expressed using an *actin (Act5C-Gal4)* driver (*BDSC– 3954: y1 w*; P{Act5C-GAL4}17bFO1/ TM6B, Tb1*), and 2) pan neuronally expressed using *embryonic lethal abnormal vision* (*elav*) driver (BDSC–25750: *P{w[+mW.hs]=GawB}elav[C155]w[1118]; P{w[+mC]=UAS-Dcr-2.D}2*). All the UAS-RNAi lines were in either P2 (Chr 3) or P40 (Chr 2) background, which we used as controls for comparing thermal tolerance.

For each line, we placed adults from the parental stock in fresh vials and allowed them to mate. On day 3, we cleared the vials of all adults, then on days 9-12, we collected 3 males (Gal4) and 5 females (UAS-RNAi) to set up the cross to obtain Gal4-UAS-RNAi F_1_ progeny. The *Act5C-Gal4* line has a tubby (*TM6B, tb1*) phenotypic marker, which we selected against at the pupae stage of development in the F1 because tubby is a pupae marker. Therefore, to make a fair comparison we also transferred all lines on days 19-22 to control for transferring the tubby lines. On days 20-23 we anesthetized the flies using CO_2_, collected 10 F_1_ females (0-1 day post-oviposition), placed them in a new vial with food, and allowed at least 12 hours of recovery time before measuring thermal tolerance.

### Measuring Thermal Tolerance in RNAi Lines

We measured the time it takes for individuals to become incapacitated in our RNAi crosses (Fig. 1C). Flies were placed individually into small clear cylinder tubes (5 mm diameter x 65 mm length; monitor tubes; TriKinetics, Inc. USA), which were plugged on both ends to prevent the flies from escaping. We placed the tubes onto a custom-milled aluminum plate (6.11mm thick) with 34 individual slots (2.1mm thick spaced 1.6mm apart) matching the diameter of the fly tubes, allowing even heating throughout the tube. The aluminum plate is mounted to a Peltier-element based heating plate (PELT-PLATE; Sable Systems International, Inc., North Las Vegas, NV), which is connected to a controller, maintaining the plate at a constant temperature of 41°C. A temperature probe was also placed into a monitor tube for each run to monitor the accuracy of the temperature within the fly tubes relative to the PELT-PLATE.

We measured each cross in three batches of ∼10 individuals per run for a total of 10 min. at 41°C. After some samples dropped out, we had measurements for between 21 and 60 individuals for each cross. Before each trial, we confirmed that the aluminum plate was at ambient temperature before placing the monitor tubes into the wells. We randomized the assay order for the lines for the first replicate and kept that order for the second and third batches.

During each 10–minute thermal tolerance trial, the entire set of flies was recorded using a Raspberry Pi High Quality Camera (12.3 MP) at 1 frame per second using a Raspberry Pi 4 Model B and custom Python code. Simultaneously, this script also recorded the timestamp, temperature reported by the PELT-PLATE, and temperature in the monitor tube measured via a thermocouple. Because the Raspberry Pi camera produces a highly distorted “fisheye” image, we first de-warped the individual images via checkerboard calibration using 15 images covering the entire plate surface (Fig. S3). The images were subsequently combined into a single movie for later analysis, both using routines in the OpenCV library (Bradski 2000).

Movies of the thermal tolerance trials were analyzed using DeepLabCut (version 2.2.3; (Mathis *et al*. 2018; Nath *et al*. 2019)) and custom Python scripts to enable batch processing of trials. Because each trial recorded the activity of many flies simultaneously, we needed to first split each composite video into a movie of a single fly’s movements. The rigid nature of the aluminum thermal plate allowed for the automated extraction of video for individual flies. First, we located the four corners of the plate using a trained DeepLabCut model (Fig. S4A). With coordinates of the four corners, we applied a second dewarping step, rotated the image so that the edges were normal to the frame, and cropped the image to the four corners using routines in the OpenCV library (Bradski 2000). Second, we leveraged the fixed coordinates of the fly tubes within the plate to subset each video into 34 separate videos, each with a single fly, no fly, or the thermocouple (Fig. S4B). These videos were then processed using a second DeepLabCut model, which had been trained on 978 manually labeled frames from individual fly movies (1,000,000 training iterations; train error = 1.64 pixels; test error = 1.69 pixels; Fig. S4C). This automated system was able to track 1,350 flies across ∼800,000 individual frames, yielding cartesian (x, y) coordinates for each fly at each 1 s timestep. With these time and position data, we used an R script to automate identifying when individual flies were incapacitated using the same criteria described for the heat box.

To determine the effect of each RNAi cross on the incapacitation phenotype, we compared the UAS-Gal4-RNAi crosses and the background control cross by fitting separate ANOVAs for each RNAi line with the cross id as the predictor and incapacitation time as the response variable. We followed this model with planned post-hoc comparisons using the *glht* function in the *multcomp* package (Hothorn *et al*. 2008), comparing each RNAi cross with the relevant matching background cross.

### Validation of Thermal Tolerance Phenotyping

Our mapping experiment used the heat box (see “Measuring thermal tolerance in the DSPR” section above) to track activity in individual flies in chambers lined individually with Peltier elements. We developed an alternative method for measuring incapacitation following this experiment for two reasons: 1) the heat box showed more variability than desired in the target temperature reached, and we identified instances where individual heat box chambers did not reach the desired temperature (these data were discarded from our dataset, see above), and 2) the heat box can measure, at most, 16 individuals at one time, limiting the speed we can perform assays. The heat plate method we developed, described above, resolved both issues, increasing the throughput of the assay and allowing for consistent assay temperatures with continuous monitoring. To confirm our phenotyping methods are comparable, we measured a subpopulation of 39 RILs from those that were assayed in the heat box. We assayed these RILs in two biological replicates, assaying ∼60 individuals each time (range: 27 individuals to 72 individuals) for a set of RILs distributed across the range of thermal tolerance scores using the heat plate apparatus and methodology (Fig. S5A). We observed shorter incapacitation times overall on the heat plate, which is expected given that the heat plate showed better consistency in reaching and maintaining the target temperature of 41° C than the heat box, but confirmed the scores are correlated between the two methods (*r* = 0.36). As with the heat box, we validated our measurements by hand-scoring a set of actual videos of flies moving for incapacitation and examined a set of traced movies to confirm both the tracking performance and the programmatic scoring of incapacitation. We compared the human scores to the computer generated scores, which had an overall positive correlation (Fig. S5B; *r* = 0.47).

All data analyses were performed in R 4.2 and lower (Posit team 2023; R Core Team 2023), python 3.9.0 (Van Rossum and Drake 2009), and DeepLabCut 2.2.3 (Mathis *et al*. 2018).

### Data and Code Availability

Raw phenotypic data for measurements of thermal tolerance both in the heat box and heat plate, including the DeepLabCut models are available from Zenodo (https://zenodo.org/record/7767713). RNA-Seq data are available from the NCBI Short Read Archive (Leinonen and Sugawara 2010) under SRA accession number: (https://www.ncbi.nlm.nih.gov/sra/SRR24828389). Raw re-sequencing data of the DSPR founder lines are deposited in the NCBI SRA under accession number SRA051316 and the RIL RAD genotyping data are available under accession number SRA051306. Founder genotype assignments from the hidden Markov model are available as two data packages in R (http://FlyRILs.org/). Additionally, see King et al. (2012b; Long *et al*. 2014) and Long et al. (2014) for more details on the DSPR datasets. The complete code used to perform all analyses is available on GitHub (https://github.com/EGKingLab/LearnMemTol).

## Results

### Phenotypic Distributions

We first quantified thermal tolerance in 741 RILs (39,392 individuals) in the DSPR using the heat box (Fig. 1A). We defined incapacitation as the time it took for an individual to become unable to move, assigned an incapacitation score, then took the mean for each RIL. We found a wide range of phenotypic variation with approximately a 15-fold range from 24 seconds – 361 seconds between RILs with low and high average incapacitation times (Fig 2). We expected thermal tolerance to show high variability among RILs because the original founding parents of the RILs were collected from different continental locations (King *et al*. 2012b) with varying adaptation to different temperatures and it is also a characteristic complex trait. We estimated broad-sense heritability (H^2^) for thermal tolerance in the RILs, which was H^2^ = 0.24, 95% CI = 0.22–0.26. In addition, the effect of RIL ID on thermal tolerance was highly significant (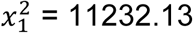, p < 0.0001), indicating that there is a genetic basis for thermal tolerance in the DSPR RILs. We also phenotype the DSPR founder lines (A1, A2, …, A7) for thermal tolerance (Fig. 3B), except forAB8, which did not thrive in our hands, and thus, we do not have an estimate for that line. We measured a total of 424 individuals, with an average of 61 individuals per founder (range: 48 – 67).

**Figure 2:**
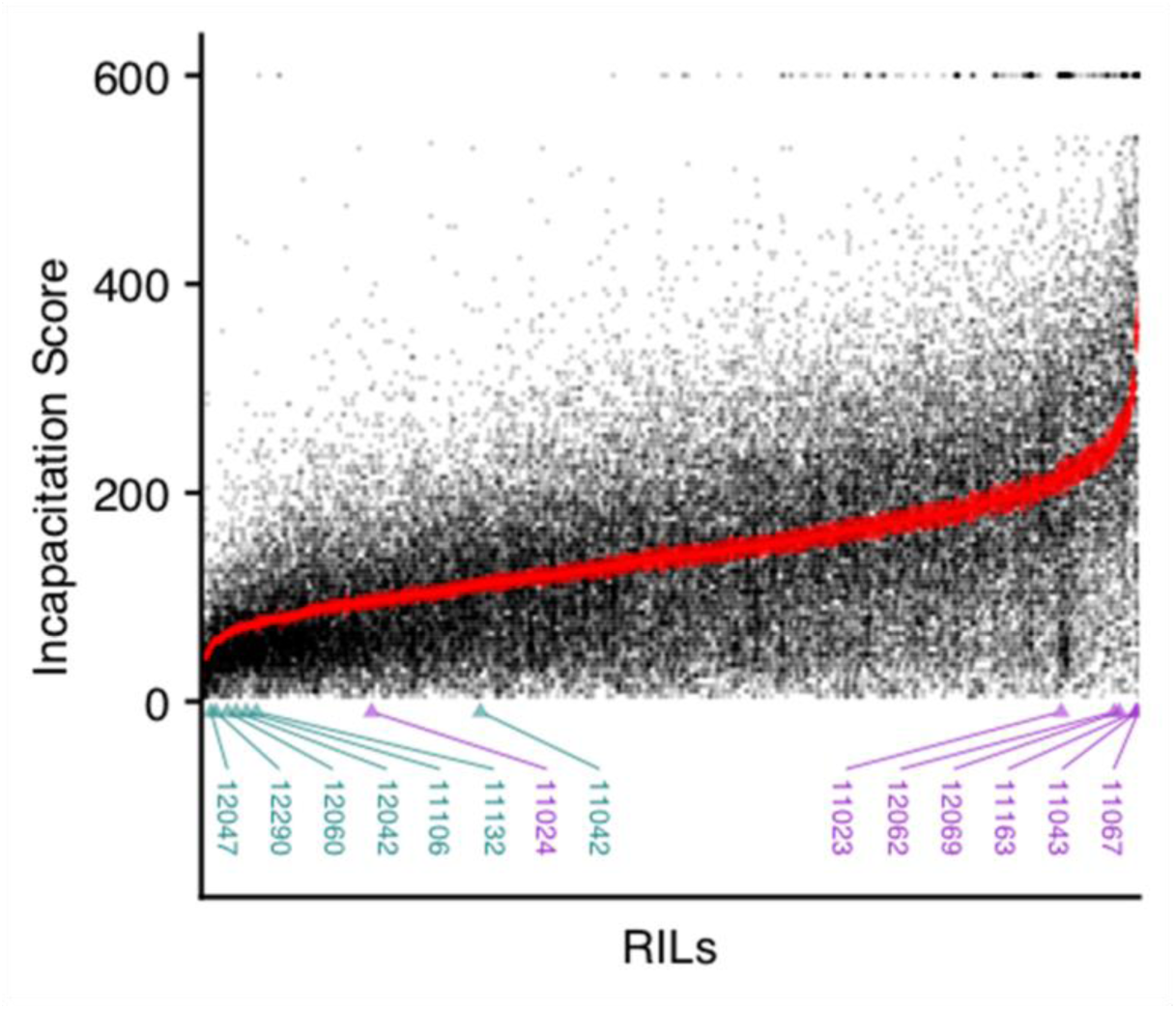
Incapacitation scores for all individuals (>48,000) for the 741 DSPR RILs measured. Each point represents an individual fly’s phenotype with points plotted with semi-transparency so overlap can be visualized as darker areas. The RILs are sorted on the x-axis by their mean incapacitation score. The mean score +/- 1 standard error for each RIL is plotted in red. The ids of the 14 RILs used for RNAseq are labeled below with the low pool RILs in dark cyan and the high pool RILs in purple.

**Figure 3:**
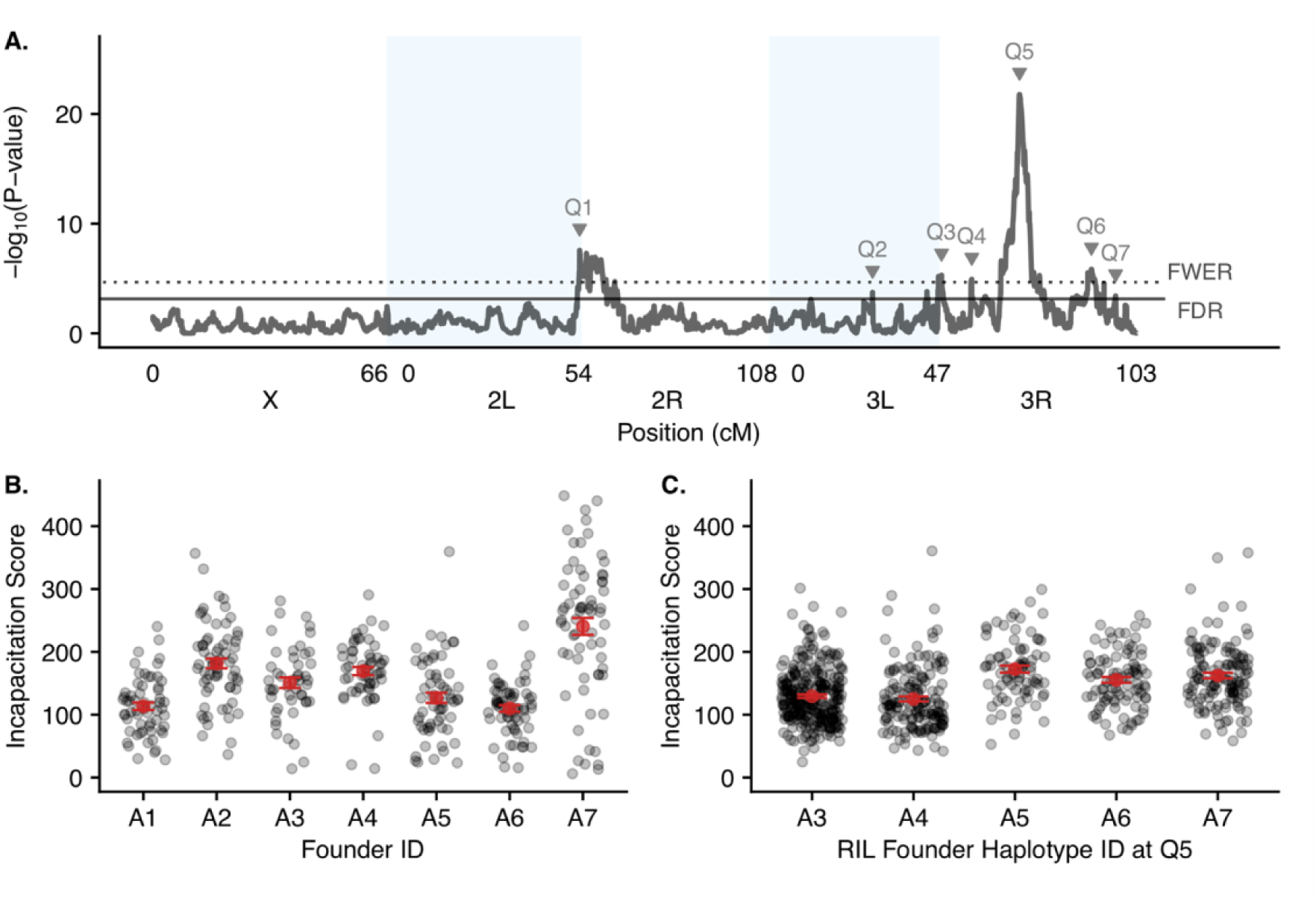
A) Genome scan for thermal tolerance in the DSPR RILs showing the -log_10_(P value) versus position in the genome with different chromosome arms denoted by shaded boxes. Each QTL that reached at least the FDR threshold is labeled at the corresponding QTL peak. The FDR (solid line) and FWER (dotted line) thresholds are shown with horizontal lines. B) The raw incapacitation scores measured in the inbred founder lines. Each point represents an individual fly’s phenotype with points plotted with semi-transparency so overlap can be visualized as darker areas. The mean score +/- 1 standard error for each founder line is plotted in red. C) The average incapacitation scores in the RILs with RILs separated by the founder haplotype they harbor at the Q6 location. Each point represents a RIL mean with points plotted with semi-transparency so overlap can be visualized as darker areas The mean score +/- 1 standard error for the set of RILs harboring each founder haplotype at Q6 is plotted in red.

### Quantitative Trait Loci

We identified 7 QTLs (Q1-Q7; Fig 3A; Table 1) at the 5% FDR and higher. Of those 7 QTLs, 5 reached or passed the FWER threshold, which is the more stringent threshold. Any significant QTLs are potential regions of the genome that influence thermal tolerance, with Q5 having the strongest association. For each significant QTL, we determined the region most likely to contain causative variants by obtaining a Bayesian Credible Interval (BCI; (Manichaikul *et al*. 2006; Broman and Sen 2009). For Q5, the most significant QTL, the BCI is 20.93 – 21.09 on chromosome 3R. An advantage of the DSPR is that we are able to infer the haplotype identity to any given location in every RIL (King *et al*. 2012b). Therefore, for each of our significant QTL, we can estimate the founder (A1, A2, A3, A4, A5, A6, A7, and AB8) haplotypes’ effect at the specific QTL location for thermal tolerance (Q5: Fig. 3C; All QTL: Fig. S6). For example, at Q5, the set of RILs harboring the A5 haplotype at that location have the highest average incapacitation score, while those harboring the A4 haplotype have the lowest, though the means do not form two distinct groups, suggesting the potential for multiple causative variants in this interval.

**Table 1:** Details for all QTLs.

| ID | Arm | Position (Mbp) | Lower (Mbp) | Upper (Mbp) | $-\log_{10}(\text{P-value})$ | PVE (%) |
| --- | --- | --- | --- | --- | --- | --- |
| Q1 | 2L | 20.11 | 2L:22.07 | 2R:8.69 | 7.57 | 6.4% |
| Q2 | 3L | 9.39 | 8.82 | 9.42 | 3.74 | 3.8% |
| Q3 | 3R | 8.02 | 5.53 | 8.28 | 5.30 | 4.9% |
| Q4 | 3R | 15.40 | 15.26 | 17.37 | 4.93 | 4.6% |
| Q5 | 3R | 20.98 | 20.93 | 21.09 | 21.79 | 14.9% |
| Q6 | 3R | 26.33 | 26.00 | 26.63 | 5.84 | 5.2% |
| Q7 | 3R | 28.94 | 26.97 | 30.40 | 3.41 | 3.6% |

### RNA-seq and Genome-Wide Expression Patterns

We performed RNA-seq on a pooled set of female heads to identify differentially expressed genes between low and high cohorts of thermal tolerance performance, assaying expression at both room temperature and following exposure to high temperature. Thus, we obtained genome-wide expression profiles (Fig. 4) for 4 separate groups: thermal tolerance high performers at room temperature (H_RT), thermal tolerance low performers at room temperature (L_RT), thermal tolerance high performers after incubation (H_Incu), and thermal tolerance low performers after incubation (L_Incu).

**Figure 4:**
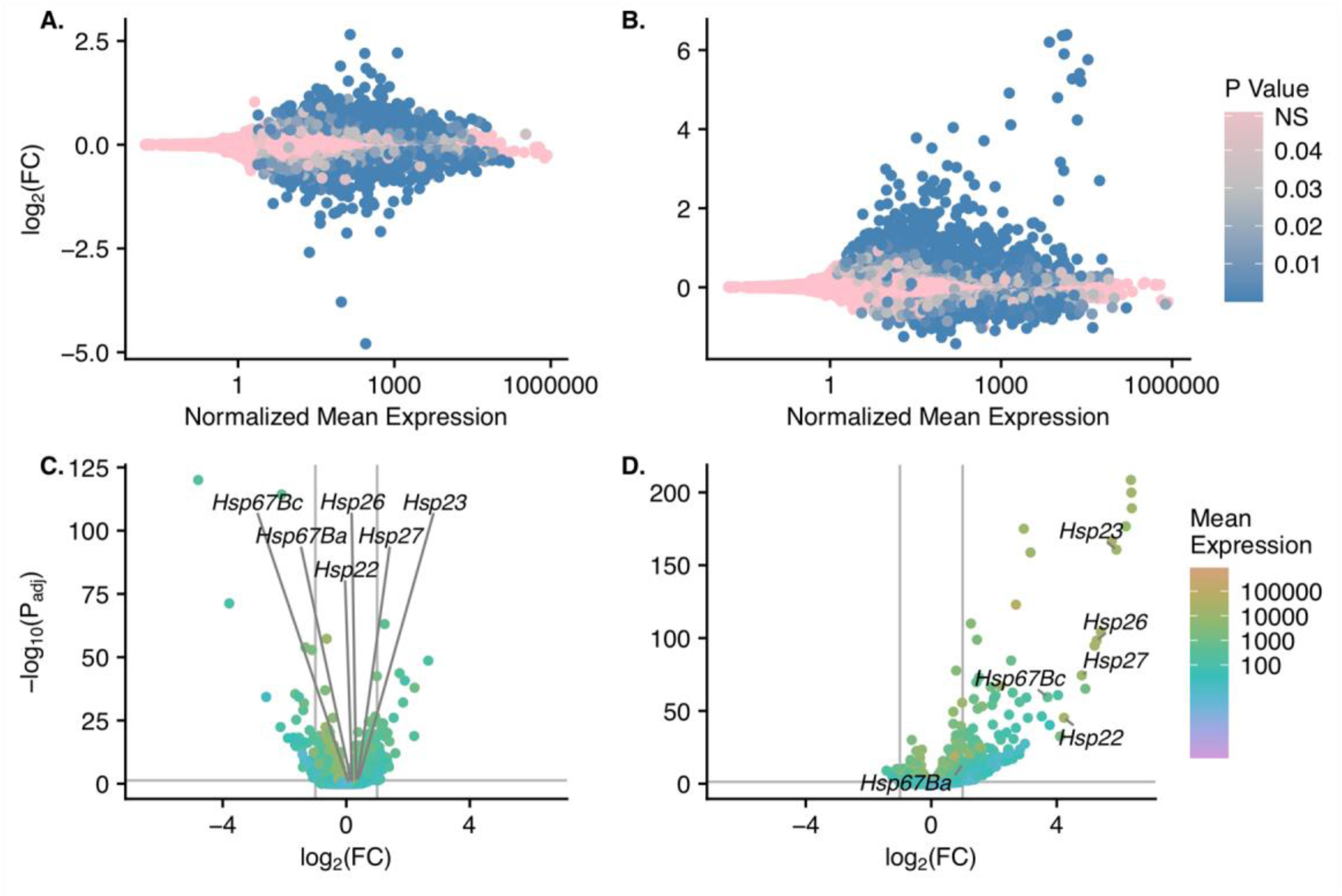
A & B) MA plots showing log_2_(fold change) for the high versus low pools (A) and the incubated versus room temperature pools (B) versus the normalized mean expression value for each gene. Normalized mean expression is shown on a log_10_ scale. Genes are colored by p-value with non-significantly differentially expressed genes in pink and increasingly significant genes in darker shades of blue. C & D) log_2_(fold change) for the high versus low pools (C) and the incubated versus room temperature pools (D) versus - log_10_(adjusted p-value). The color gradient of the points shows the normalized mean expression value of each gene. The *Hsp70* family of genes is labeled on each plot.

First, to determine the level of genome-wide expression patterns in the different cohorts, we performed principal components analysis (PCA) on all samples using the expression matrix (Fig. S7). PC1 explains 29% of the variance, while PC2 explains 22% of the variance. There is a clear separation between the samples incubated at high temperatures and those measured at room temperature. Both high and low-performance pools following high-temperature exposure (Incu) cluster together, whereas there is a substantial separation between high and low pools (H and L) at room temperature (RT).

Next, we determined differences in gene expression patterns in the different cohorts (Fig. 4). There were 2642 genes that were differentially expressed between high vs. low (pool) cohorts for thermal tolerance genome-wide. In this comparison, 1215 genes had higher expression in the high pool and 1427 had higher expression in the low pool (Fig. 4). Of those genes, 191 were located between our QTL’s BCI (Q2 = 18; Q3 = 49; Q4 = 35; Q5 = 8; Q6 = 15; Q7 = 66; Fig. S8), obviously, higher numbers of genes are located under the QTLs with wider BCIs. Q1 is excluded here because the BCI spans a centromere. In the incubated vs room temperature (condition) comparison, 2862 genome-wide genes were differentially expressed between high and low cohorts for thermal tolerance. In this comparison, 1608 genes had higher expression in the incubated condition and 1254 had higher expression at room temperature. There were 256 genes differentially expressed within our QTL intervals (Q2 = 29; Q3 = 66; Q4 = 59; Q5 = 8; Q6 = 12; Q7 = 82; Fig. S9).

We then wanted to examine if any Heat Shock Protein (HSP) genes within our QTL intervals were differentially expressed within the cohorts (pool or condition) because *Hsp*s are characterized as genes important for thermal tolerance in fruit flies and other species (Krishnan *et al*. 1989; Sørensen *et al*. 2001; Pörtner 2002; Fangue *et al*. 2006; Lockwood *et al*. 2017). Within Q2, which is located on chromosome *3L*, there were 6 differentially expressed *Hsp* genes (*Hsp22, Hsp23, Hsp26, Hsp27, Hsp67Ba,* and *Hsp67Bc*) in the condition cohort whereas only *Hsp23* was significant in the pool cohort.

### Candidate Genes in the Q5 Region

Among our identified QTLs, Q5 was much more significant than the other peaks (Fig. 3A), indicating that this locus has a larger effect on thermal tolerance, and therefore we consider Q5 a QTL of high interest. Q5 is located on chromosome 3R there are 21 genes located within that BCI region (Fig. S8D; S9D). Eight of the 21 genes were significantly differentially expressed between high and low pools at room temperature: *annexin B9 (anxB9), cortactin (cortactin), dynein heavy chain (dhc93AB), malvolio (mvl), Na pump alpha subunit (α), rho GTPase activating protein (RhoGAP93B), CG31207* and *CG7079.* We focused on differential expression based on standing expression in the low and high pools at room temperature because our thermal tolerance assay is short (10 minutes) and changes in gene expression in response to temperature would require longer timescales to impact the phenotype.

### RNAi

In order to investigate the effect of differences in gene expression for genes within the Q5 interval on thermal tolerance we used the TRiP lines with knock-down expression of a particular gene. We quantified thermal tolerance in 20 UAS-RNAi mutant lines (1969 individuals), representing 14 genes (Table S2), which we crossed to Act-Gal4 (all cells) and elav-Gal4 (all neurons). We observed significant phenotypic effects on several of these crosses (Fig. 5; Fig. S10; Table S2), with the most common effect being a decrease in incapacitation time compared to the background cross. Adding this RNAi evidence to our existing data, we selected five genes (*anxB9, cortactin, Dhc93AB, Mvl, and CG31207*) as possible candidates, based on several lines of evidence.

**Figure 5:**
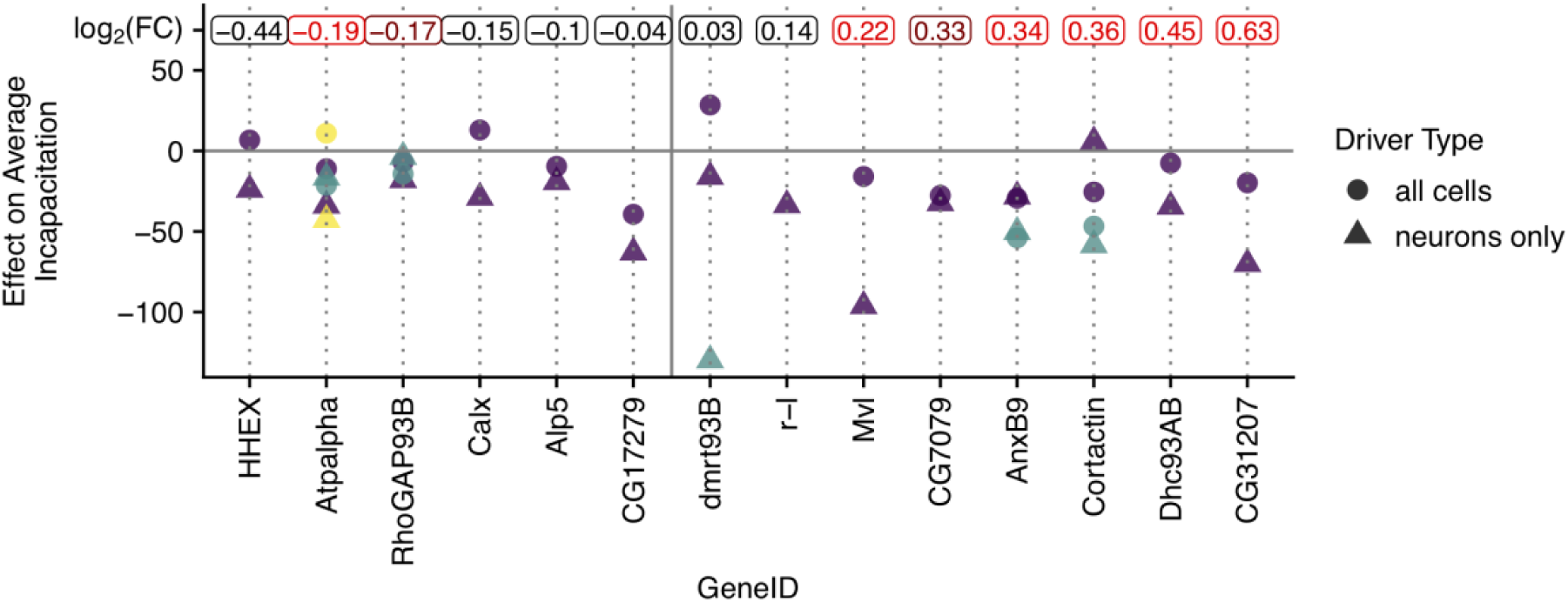
The effect of RNAi knockdown for each assayed gene within the Q6 BCI. Plotted points show the difference between average incapacitation scores between each RNAi cross and the corresponding background cross for each gene within the Q6 BCI assayed. Negative numbers indicate the RNAi cross had a lower average incapacitation score than the matching background cross. Different shapes show drivers targeting different sets of tissues with circles representing the driver targeting all cells (ubiquitously expressed) and triangles showing the driver specific to neurons (pan-neuronal). Some genes had more than one RNAi line available so for those cases, different colors represent the different RNAi lines. In some cases, all crosses for a particular driver - RNAi line combination was not successful and thus those are not plotted. Genes are ordered on the x axis by the degree of differential expression between pools of lines showing high versus low incapacitation. The log_2_(fold change) for each gene is displayed at the top of the plot with significantly differentially expressed genes shown in red (dark red = 0.01 < p < 0.05; red = p < 0.01).

First, we started with a list of all the genes that were located within the BCI for Q5 and that were also significantly differentially expressed in our RNA-seq dataset (Fig. 5; Fig. S11). Second, we combined this list with the RNAi data, where we measured the changes in incapacitation compared to the background line when that particular gene was knocked down. We considered a gene worthy of further consideration if the direction of the RNAi effect was consistent with our RNAseq results (e.g., a gene with higher expression in the high thermal tolerance group should show a decrease in thermal tolerance in the RNAi cross), this effect was significant for at least one cross for that focal gene, and there was overall consistency in the direction of the effect in the set of crosses for the gene. The one exception to this criterion is *Dhc93AB*, which we include because it was one of our most significantly differentially expressed genes and the phenotypic effect of the RNAi line was consistent with the expected effect, though this effect was not significant. The following genes met these criteria: *anxB9, cortactin, Dhc93AB, Mvl, and CG31207,* and the effects and significance for these genes are shown in Table 2. While not every RNAi cross associated with a given gene showed a statistically significant effect, there is general consistency in the direction of the effect for the genes in our candidate list across crosses.

**Table 2:**
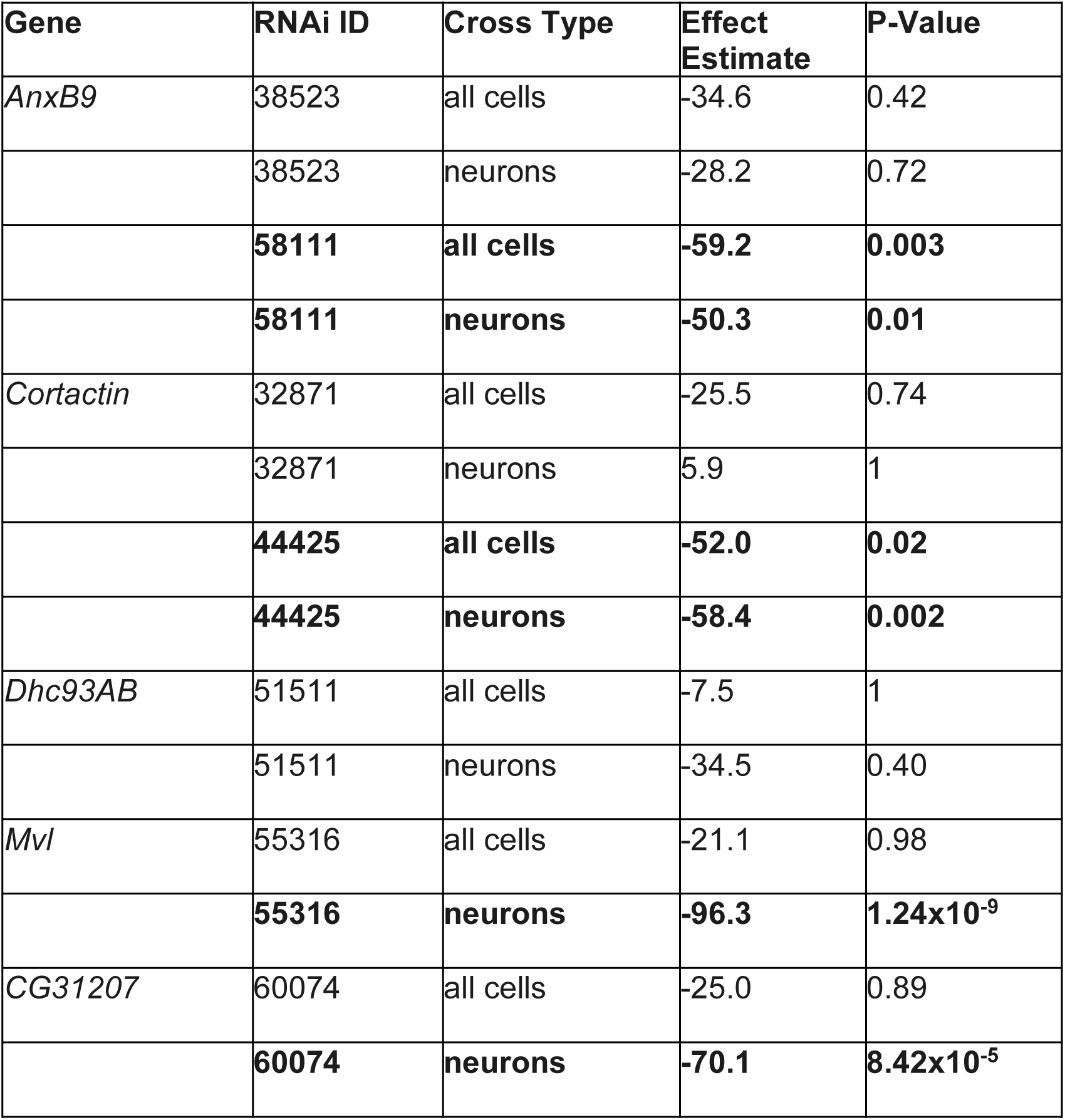
Effect estimates and P-Values from post-hoc tests for each RNAi cross in our candidate list.

| Gene | RNAi ID | Cross Type | Effect Estimate | P-Value |
| --- | --- | --- | --- | --- |
| *AnxB9* | 38523 | all cells | -34.6 | 0.42 |
|  | 38523 | neurons | -28.2 | 0.72 |
|  | **58111** | **all cells** | **-59.2** | **0.003** |
|  | **58111** | **neurons** | **-50.3** | **0.01** |
| *Cortactin* | 32871 | all cells | -25.5 | 0.74 |
|  | 32871 | neurons | 5.9 | 1 |
|  | **44425** | **all cells** | **-52.0** | **0.02** |
|  | **44425** | **neurons** | **-58.4** | **0.002** |
| *Dhc93AB* | 51511 | all cells | -7.5 | 1 |
|  | 51511 | neurons | -34.5 | 0.40 |
| *Mvl* | 55316 | all cells | -21.1 | 0.98 |
|  | **55316** | **neurons** | **-96.3** | **1.24x10<sup>-9</sup>** |
| *CG31207* | 60074 | all cells | -25.0 | 0.89 |
|  | **60074** | **neurons** | **-70.1** | **8.42x10<sup>-5</sup>** |

Furthermore, we investigated if there were any structural variants located within the DSPR founders with haplotypes located at the Q5 region, following the protocol of (Chakraborty *et al*. 2019). Structural variants are hidden mutations within the genome that are known to shape variation in complex traits (Fraser and Xie 2009; Eichler *et al*. 2010; Chakraborty *et al*. 2019), and therefore identifying them is a critical step in dissecting complex traits. We found four structural variants that fall within Q5 (Table S4), including an insertion within *anxB9,* that occurs in the founder A3 haplotype. Interestingly, the A3 founder has one of the lower incapacitation scores (Fig 3C) at Q5. Within all 7 QTLs, we found a total of 413 SVs (Table S3). Lastly, we performed a manual search on FlyBase (Gramates *et al*. 2022) to investigate previous findings of these genes being implicated with thermal tolerance. None of the five genes have been shown to have direct implications for thermal tolerance previously. Thus, our results showing both differential expression in low and high-performing cohorts and a consistent effect following RNAi knockdown of these same genes within a large effect QTL are the first suggestion that these genes may be involved in thermal tolerance.

## Discussion

We identified 7 genomic regions that influence thermal tolerance in the DSPR, including one large effect locus on chromosome *3R*. We also identified multiple differentially expressed genes between cohorts that differ in high vs. low thermal tolerance. For our most significant QTL, we integrated our RNA-seq findings and RNAi results to identify several candidate genes with multiple lines of evidence for their involvement in thermal tolerance. Overall, we found evidence for multiple potential causative variants within the large effect locus on Q5, supporting the hypothesis that in some cases, mapped QTLs do not represent a single causative variant, but instead a collection of variants acting in concert.

Our results expand on previous studies that took genome-wide approaches to investigate the genetic basis of thermal tolerance in *D. melanogaster (Norry et al. 2004, 2007a; b, 2008; Morgan and Mackay 2006; Rand et al. 2010; Rolandi et al. 2018)*. There are multiple different methods for measuring thermal tolerance, and thus there are some limits to direct comparisons between our study and previous work. Nevertheless, it is essential to observe if there might be common loci identified in these different studies. We first compared our QTL dataset to Norry et al. (2008), because they characterized thermal tolerance in two divergent populations that were measured for thermal tolerance in both males and females. Norry et. al (2008) used heat-hardening measurements (exposure to high temps repeatedly) with an exposure temperature similar to our measurements. We found that our Q2 had an overlap with the data that was presented in that study, however, when we compared to the list of candidate genes only *Hsp23* was common among the two groups. This is not surprising, because, Frydenberg et al. (2003), performed genomic analysis on isofemale lines with a specific focus on *Hsp23*, *Hsp26*, and *Hsp27*, and found that at *Hsp23,* allele frequency differs across latitude clines in *D. melanogaster*. Interestingly, both the founding parents from DSPR – which we used in this study – and the RILs used in Norry et al., (2008) were collected in different latitudinal climatic locations, and represent different sets of genotypes, which could also contribute to the differences in the studies. We also compared our study to the findings of Rolandi et al. (2018). This group studied the genetic basis of thermal tolerance by measuring the upper thermal limits of 34 lines from the DGRP and performed GWAS to identify QTLs. The DGRP – like the DSPR – is one of the multi-parental populations in fruit flies. Briefly, they reported 8 major SNPs that are associated with upper thermal limits in both males and females. We did not find any overlap between the SNPs they identified as candidates and our QTLs, nor did they correspond with genes that were significantly expressed within our QTLs. We also cross-referenced the candidate genes found in Norry et al., (2008) and Rolandi et al. (2018) to our list of genes that were significant between high and low thermal tolerance pools and found only one overlap – *Hsp23* (Norry *et al*. 2008). We have not directly compared effect estimates for the loci identified in these previous studies and our own, thus, loci we did not identify in this study may still be consistent with those findings and not identified in our study due to power. Thus, while this comparison to previous work does not allow us to make conclusions about differences in the genetic basis of thermal tolerance in these different study populations, this comparison does show that we have identified a set of novel QTLs and candidate genes that potentially influence thermal tolerance in the *D. melanogaster,* adding to our knowledge about the genetic basis of this critical trait.

Our findings emphasize the natural phenotypic and genetic variation in thermal tolerance (Sørensen *et al*. 2001, 2019; Norry *et al*. 2004, 2008; Overgaard and Sørensen 2008; Overgaard *et al*. 2011; Rolandi *et al*. 2018; MacLean *et al*. 2019a; b). This approach contrasts with a mutagenesis approach, which, while powerful, does not focus on natural genetic variation that is segregating. For example, we did not identify any *Hsp70* genes in our integrated approach comparing high and low thermal tolerance groups. This gene family has been well-established as having a role in heat tolerance, which stems from much of the earlier work done on elucidating mechanisms of thermal tolerance (Welte *et al*. 1993; Feder *et al*. 1996). However, more recent work by Jensen et al. (2010) suggests *Hsp70* is not related to heat tolerance variation in adult *D. melanogaster.* Additionally, a GWAS study by Lecheta et al., (2020), also did not identify any *hsp* genes in their findings. The only gene from the *Hsp* family that was significantly differentially expressed within our design is *Hsp23*, within our pool treatment. Another potential reason for the lack of *Hsp70* genes in our study could be that most of the studies that identified *Hsp70* using *D. melanogaster* had a longer exposure or repeated exposure (Norry *et al*. 2008; Rampino *et al*. 2009; Cleves *et al*. 2020), whereas our measurement of thermal tolerance was a total of 10 minutes. Our focus on natural genetic variation is one of the advantages of the DSPR. In the DSPR system, the fly genome is unmodified (natural), however, we can rear them in the lab within a controlled environment, therefore reducing possible uncontrollable confounding effects. This approach allows for unbiased genomic scans to uncover natural segregating variants that would likely not be identified using single-gene approaches. Because of this systematic approach, we can identify multiple genome loci that influence complex traits, like thermal tolerance.

Genome scans are often paired with RNA-seq to further dissect the genetic basis of complex traits. We integrated RNA-seq to identify differentially expressed genes between high and low thermal tolerance cohorts within our 7 QTL regions (Wayne and McIntyre 2002). This integration continues to be a powerful approach used to narrow down potential candidate genes within QTLs that affect complex traits. We compared both low and high (baseline at room temp) groups and incubated vs. room temp for genome-wide expression. This approach allowed us to identify a large set of genes whose expression differs constitutively among high and low thermal tolerant lines and the set of genes with expression changes following exposure to high temperatures. This rich dataset complements our QTL mapping results, giving additional information about the connection between gene expression and thermal tolerance. Often, the resolution of mapping approaches presents a challenge, with tens to hundreds of genes located within an associated QTL. For example, there are 472 genes located within the BCI of Q7 for the high and low cohorts, however, through RNA-seq, we were able to narrow down this list to 66 candidate genes. Notably, we might be excluding potential coding variants with this approach, however, initially focusing on regulatory variants was the most feasible approach in this case. An important limitation of our study is that we selected the 7 high and 7 low RILs for RNA-seq after initially measuring 140 RILs. Unfortunately, after measuring additional individuals and an additional 601 RILs, RILs in the high and low cohorts do not completely represent the tails of the thermal tolerance distribution. Despite this limitation, the high and low pools remain enriched for high versus low thermal tolerance lines (Fig. 1a).

To functionally validate the QTL, we used the TRIP resource (Perkins *et al*. 2015), which is a powerful resource that provides the ability to screen more than 71% of the genes in the fly genome, because there is a mutant line for almost all the genes. We took advantage of this resource and ordered lines with gene disruption for the genes that fall within our Q5 region. In total, we screened 20 lines that we crossed to two pan-neuronal (*elavGAL4*) fly lines and one ubiquitously expressed line (*act-Gal4*). We then measured thermal tolerance in those lines together with background controls to identify how the reduction in the expression of our candidate genes influenced thermal tolerance. As hypothesized, we were not able to identify a single causal gene for thermal tolerance, but instead, our results are consistent with several genes contributing to the phenotype (Fig 5). In fact, recent work by Herrmann and Yampolsky (2021) suggests that approximately 2000 genes (⅛ of the Drosophila genome) either directly or indirectly contribute to temperature adaptation, suggesting a highly polygenic architecture with most variants having small effects.

Here, we present evidence for some of the genes we found to be influencing thermal tolerance, though the other genes within the QTL interval should not be discounted at this point. We highlight five lines that we consider to show interesting patterns. Those lines had a knockdown expression of *anxB9, cortactin, Dhc93AB, Mvl, and CG31207*. Based on a FlyBase and brief literature search, it is worth noting that none of these five genes have been previously shown to directly influence thermal tolerance. In fact, relatively little is known about these genes. Briefly, *anxB9* is shown to be involved in the biological processes described in transportation, localization, and development, which is concluded based on work done mostly in mutants (Bejarano *et al*. 2008; Tjota *et al*. 2011). *Cortactin* is involved in cell organization and reproduction, in addition, to transport and development (Somogyi and Rørth 2004; Quinones *et al*. 2010; O’Connell *et al*. 2019), while *dhc93AB* is involved in nervous system processes, specifically sensory perception and movement (Gaudet *et al*. 2011; Senthilan *et al*. 2012). The gene *mvl* is shown to be involved in other nervous system processes and respond to stimuli, which were also inferred from *D. melanogaster* mutants (Orgad *et al*. 1998; Southon *et al*. 2008; Panda *et al*. 2013). Very little is known about *CG31207*, and thus our work presents some of the first phenotypic data to be associated with this gene.

Our study shows the challenges of identifying the genetic basis of highly polygenic traits even with a high powered design, an advanced QTL mapping panel, and multiple informative datasets. However, we have also demonstrated how we can gain insightful information from a multi-pronged approach, even if we do not identify “a single gene” associated with a particular QTL. Instead, taken together, our results are consistent with the hypothesis that there are multiple contributing variants within the large effect QTL we identify and additional contributing loci across the genome. Overall, our findings suggest a highly polygenic architecture for thermal tolerance, which may be characteristic of many complex traits, as has been suggested previously (Rockman 2012). This model of the genetic architecture of complex traits has a long history, from the infinitesimal model proposed by Fisher (1918), to the more recent omnigenic model developed from this foundation (Boyle *et al*. 2017). Future approaches to the investigation of the genetic mechanisms underlying thermal tolerance should recognize this inherent complexity and consider pulling in additional intermediate phenotypes and methods for assaying heat tolerance.

## Acknowledgments

This project was only possible due to the leadership and enthusiasm of Dr. Troy Zars, who sadly passed away in 2017. We thank the undergraduate researchers in Troy Zars lab, namely James Mrkvicka, Samuel Mitchell, and Christopher Posey, who helped with collecting preliminary data for this project. We also thank Dr. Stuart Macdonald for supplying us with the DSPR RILs. Rapid Genomics (Gainesville, FL) conducted the RNA extractions, library prep, and readouts, and the University of Missouri Research Computing Support Services, in particular Christopher Bottoms and Jacob Gotberg, for their bioinformatics computing support. We also thank Paul Schmidt for comments on helping to improve this paper. In addition, we thank Jacelyn Shu (https://www.jacelyndesigns.com) for the experimental design figure. This material is based upon work supported by the National Science Foundation under Grant No. IOS-1654866 (T.Z. and E.G.K.), National Institutes of Health R01GM117135 (E.G.K.), MU Research Council Grant URC-16-007 (T.Z.), National Institute of Health IMSD R25GM056901 (P.A.W-S.), and an HHMI Gilliam Fellowship (P.A.W-S.).

