## Supplementary Information for "Naturally segregating genetic variants contribute to thermal tolerance in a *D. melanogaste*r model system"

**Supplementary Tables**

Table S1: Fly food diet components

| | |
|---|---|
| Water (DI) | 1L |
| Dextrose | 129.4g |
| Agar | 6.25g |
| Yeast | 32.4g |
| Yellow cornmeal | 61.2g |
| Tegosept | 2.7g |
| 95% EtOH | 11mL |

Table S2: Post-hoc tests for each RNAi cross

| Gene | RNAi ID | Cross Type | Effect Estimate | P-Value |
|---|---|---|---|---|
| *Alp5* | 57526 | all cells | -14.8 | 1 |
| | 57526 | neurons | -19.3 | 0.99 |
| *AnxB9* | 38523 | all cells | -34.6 | 0.42 |
| | 38523 | neurons | -28.2 | 0.72 |
| | **58111** | **all cells** | **-59.2** | **0.003** |
| | **58111** | **neurons** | **-50.3** | **0.01** |
| *Atpalpha* | 32913 | all cells | -11.1 | 1 |
| | 32913 | neurons | -33.8 | 0.74 |
| | 33646 | all cells | -21.4 | 0.91 |
| | 33646 | neurons | -16.5 | 1 |
| | 51411 | all cells | 5.7 | 1 |
| | **51411** | **neurons** | **-42.7** | **0.03** |
| *Calx* | 28306 | all cells | 13.0 | 1 |
| | 28306 | neurons | -28.9 | 0.70 |
| *Cortactin* | 32871 | all cells | -25.5 | 0.74 |
| | 32871 | neurons | 5.9 | 1 |
| | **44425** | **all cells** | **-52.0** | **0.02** |
| | **44425** | **neurons** | **-58.4** | **0.002** |
| *Dhc93AB* | 51511 | all cells | -7.5 | 1 |
| | 51511 | neurons | -34.5 | 0.40 |
| *dmrt93B* | 27657 | all cells | 28.6 | 0.56 |
| | 27657 | neurons | -16.0 | 1 |
| | **50656** | **neurons** | **-129.9** | **$<1x10^{-16}$** |
| *HHEX* | 60119 | all cells | 1.5 | 1 |
| | 60119 | neurons | -23.9 | 0.90 |

| | | | | |
|---|---|---|---|---|
| *Mvl* | 55316 | all cells | -21.1 | 0.98 |
| | **55316** | **neurons** | **-96.3** | **$1.24x10^{-9}$** |
| *r-l* | 55183 | neurons | -33.6 | 0.42 |
| *RhoGAP93B* | 31167 | all cells | -6.9 | 1 |
| | 31167 | neurons | -18.0 | 0.99 |
| | 35027 | all cells | -14.3 | 0.99 |
| | 35027 | neurons | -3.6 | 1 |
| *CG7079* | 61174 | all cells | -32.9 | 0.51 |
| | 61174 | neurons | -32.3 | 0.49 |
| *CG17279* | 60071 | all cells | -44.6 | 0.09 |
| | **60071** | **neurons** | **-63.0** | **$7.45x10^{-4}$** |
| *CG31207* | 60074 | all cells | -25.0 | 0.89 |
| | **60074** | **neurons** | **-70.1** | **$8.42x10^{-5}$** |

Table S3: Number of structural variants within each QTL BCI.

| Location | Number of SVs |
|---|---|
| Q2 | 16 |
| Q3 | 0 |
| Q4 | 111 |
| Q5 | 4 |
| Q6 | 45 |
| Q7 | 237 |

Table S4: List of structural variants that are located within Q5 BCI.

| Location | Gene Name | Type of SV | Founder | SV ID |
|---|---|---|---|---|
| 3R:20947530-20947530 | NA | Insertion | A3 | 0000008965 |
| 3R:20997328-20997362 | *calx* | Insertion | A6 | 0000011920 |
| 3R:21067722-21067722 | *anxB9* | Insertion | A3 | 0000008970 |
| 3R:21074479-21074712 | *dmrt93B* | Deletion | A5 | 0000017347 |

**Supplementary Figures**

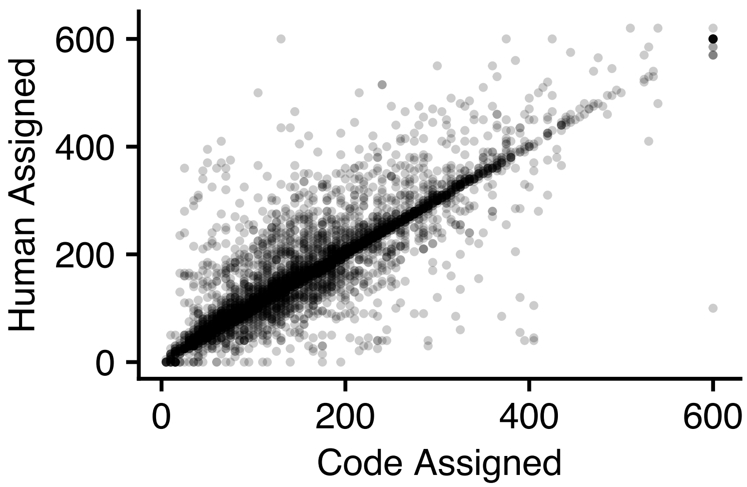


Figure S1: Human assigned incapacitation times versus automated incapacitation values via a custom R function.

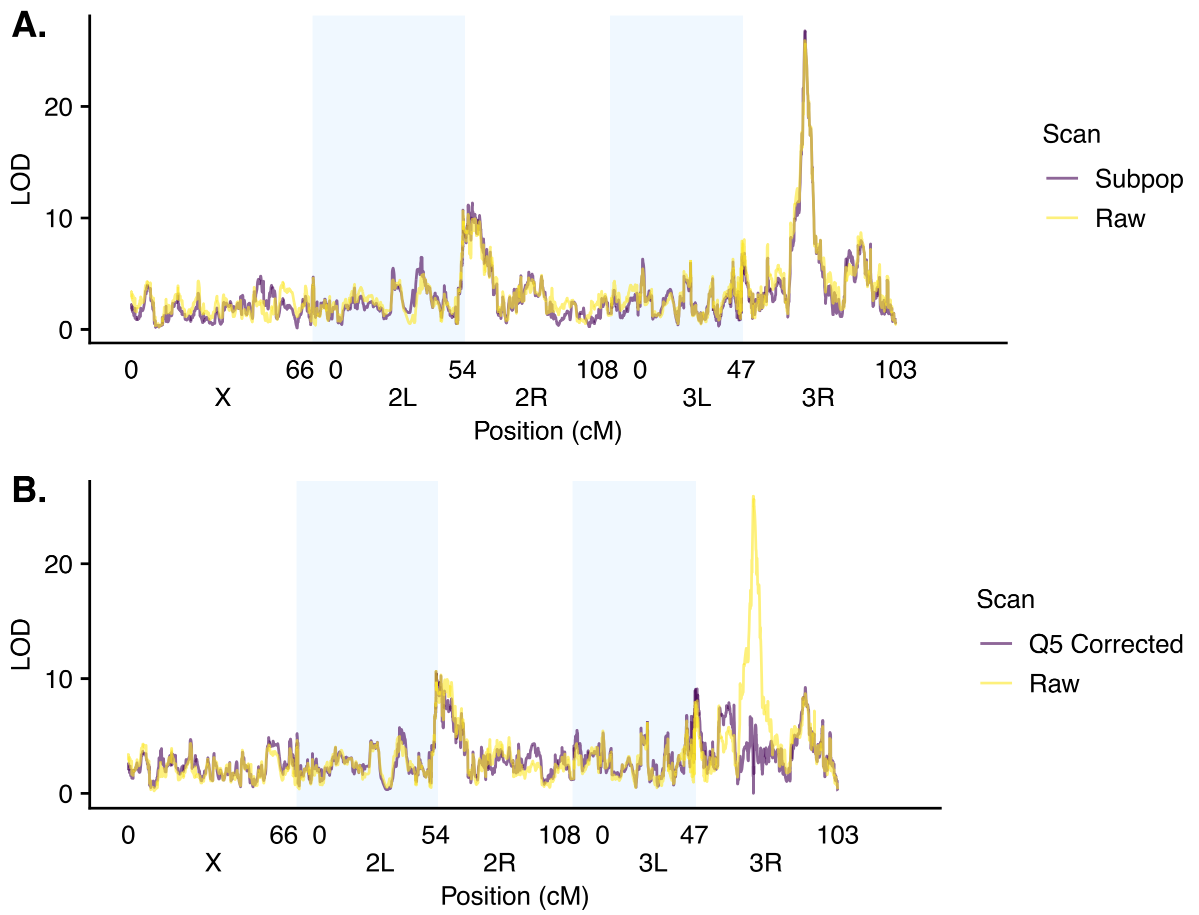


Figure S2: A) Comparison of a genome scan for a model without subpopulation (Raw, yellow line) and a genome scan for a model that includes subpopulation (Subpop, purple line). B) Comparison of a genome scan for a model without correcting for the large Q5 peak (Raw, yellow line), and a genome scan that corrects for the Q5 peak (Q5 Corrected, purple line) by including the haplotype probabilities at the Q5 location in the model. Different chromosome arms are denoted by shaded blue boxes.

A.

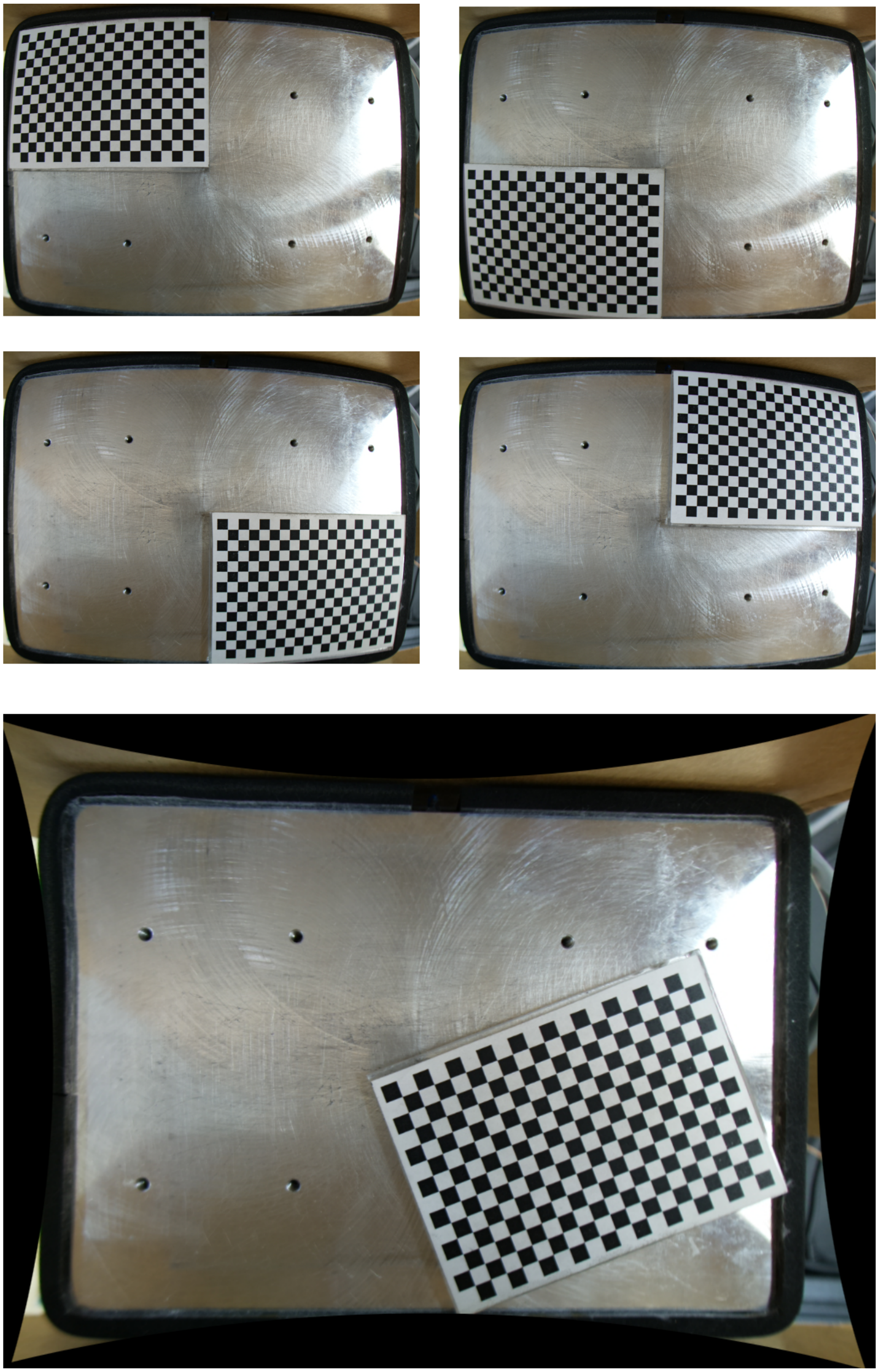

B.

Figure S3: The image dewarping process. A) Select original images (4 of 15) from the Raspberry pi camera with the calibration grid in different positions. B) The dewarped image after calibration showing the same calibration grid as in (A).

A.
B.
C.

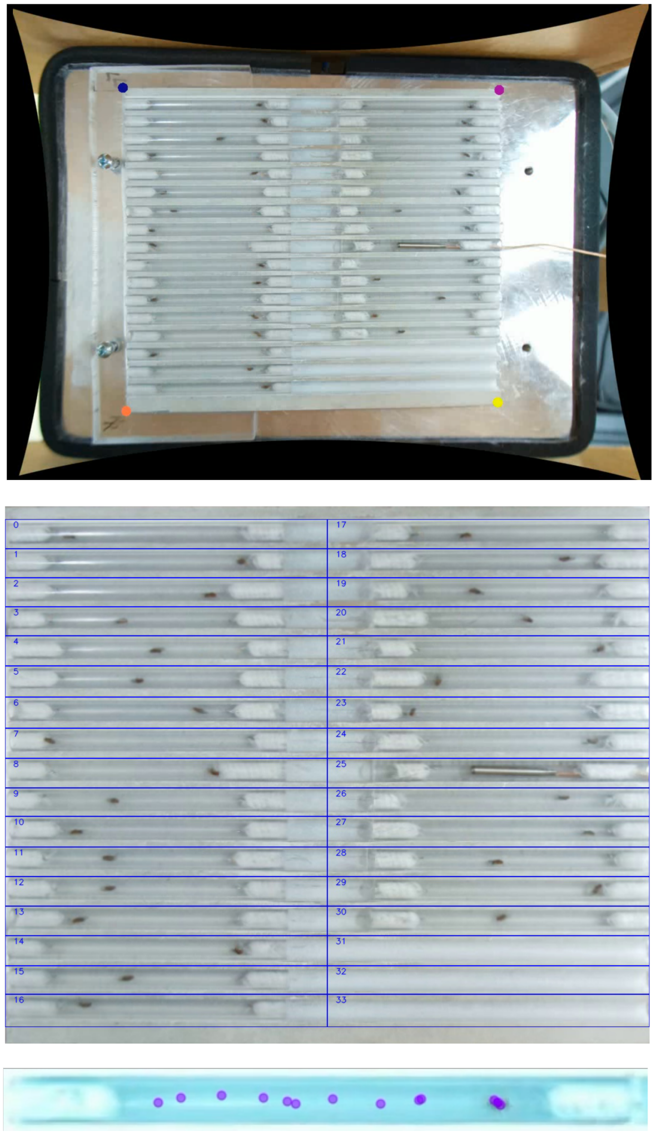


Figure S4: The image processing process to track fly activity. A) The heat plate with the aluminum holder filled with individual tubes each containing a single fly. The corners are identified automatically using a trained DeepLabCut model and are marked with large colored points. B) After squaring and cropping, the resulting grid marking each individual fly tube allowing for each video to be separated and individual flies to be tracked. C) An individual fly tube with points showing how the trained DeepLabCut model tracks an individual fly's position.

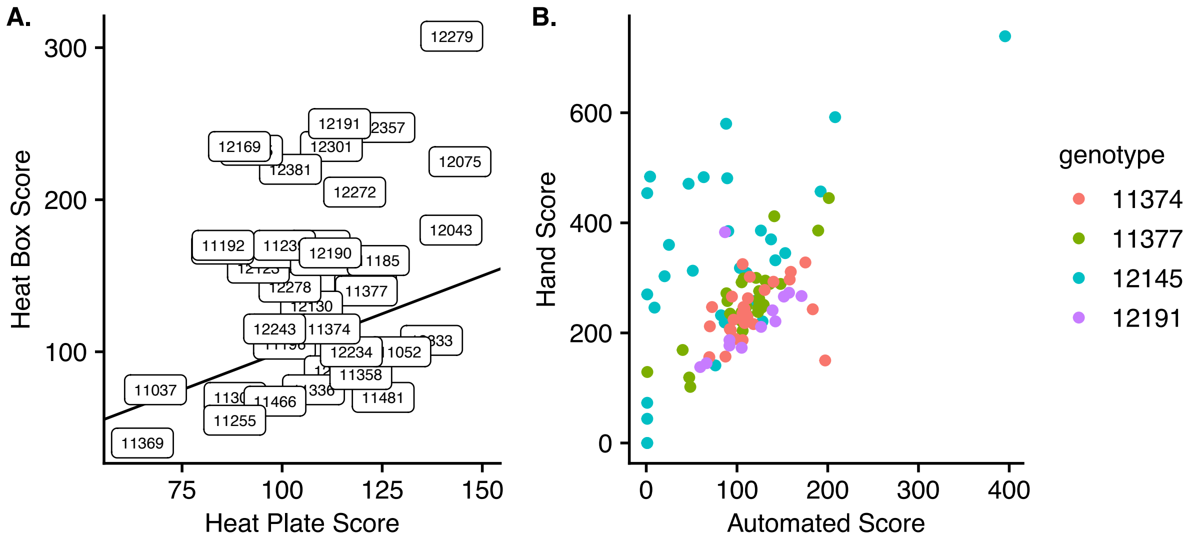


Figure S5: A) Incapacitation scores for a set of RILs measured in the heat plate apparatus versus the same RILs measured in the heat box apparatus. Each RIL id is plotted showing the scores in each. The 1:1 line is also plotted for context. B) Automated incapacitation scores from a custom R script versus hand scores for a set of four RILs measured in the heat plate. Note that the videos used for hand scoring included all frames but when the framerate exceeded 1 per second, the time on the video is inflated so those scores are higher overall.

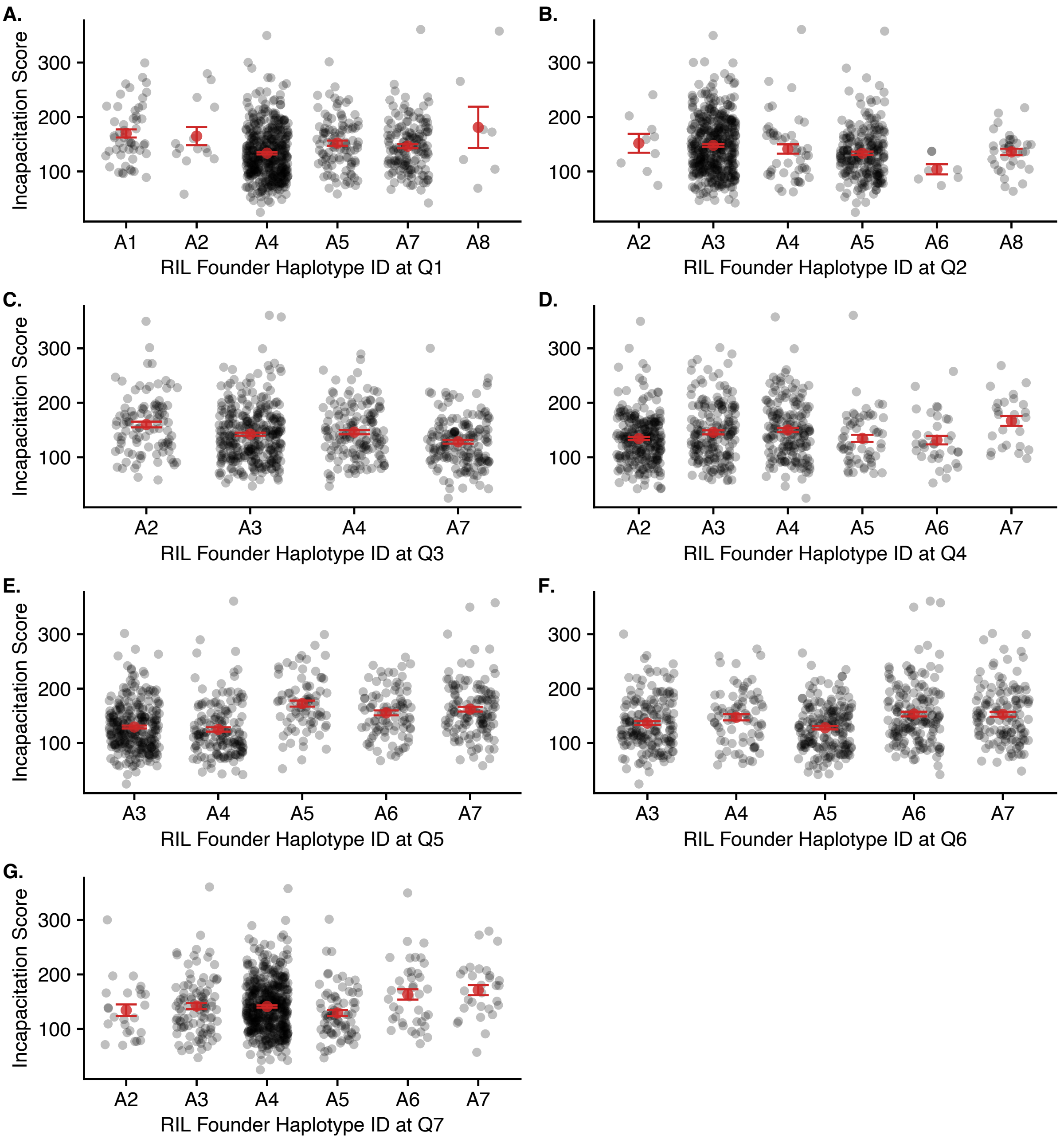


Figure S6: The average incapacitation scores in the RILs with RILs separated by the founder haplotype they harbor at each QTL location (A - H = Q1 - Q8). Each point represents a RIL mean with points plotted with semi-transparency so overlap can be visualized as darker areas. Only founder haplotypes for which there are at least 4 RILs with that haplotype at the QTL are plotted. The mean score +/- 1 standard error for the set of RILs harboring each founder haplotype at the relevant QTL is plotted in red.

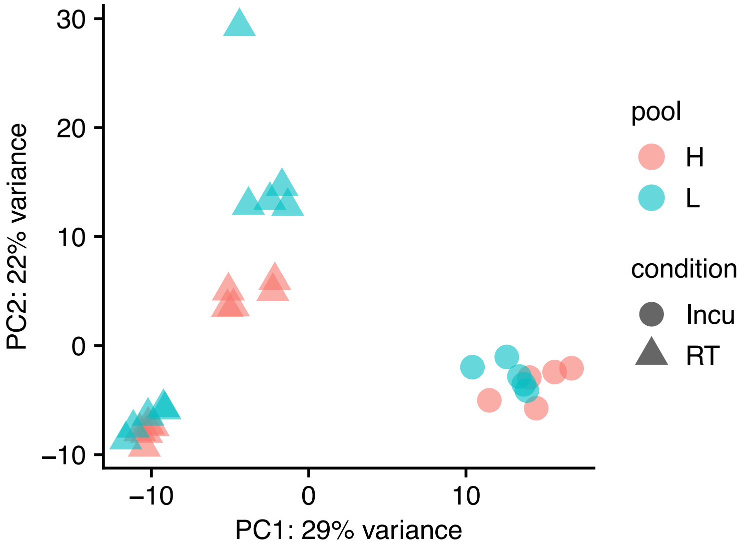


Figure S7: Principal component analysis (PCA) showing the global effect of temperature on high and low cohorts. Different colors indicate the different pools of RILs with the pool of high thermal tolerance RILs in red and low in blue. Shape denotes different temperature environments (incubated at 41°C = Incu (circles) and room temperature at 24°C = RT (triangles)). Two PC dimensions are shown here: PC1 on the x-axis and PC2 on the y-axis.

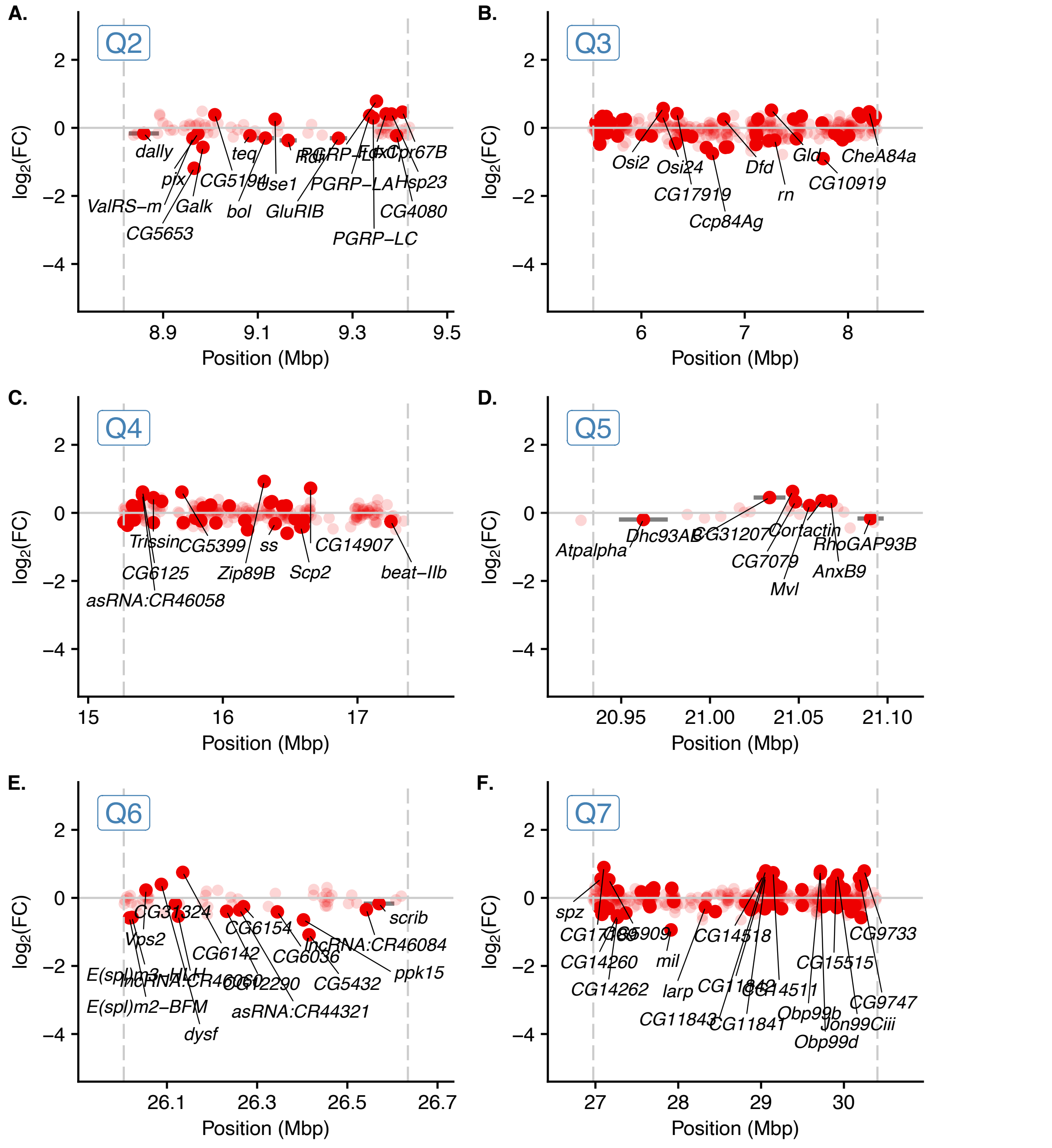


Figure S8: The $\log_2$(fold change) between expression levels in high and low performance thermal tolerance pools for each gene within each QTL BCI region. Each QTL id is labeled on the top left of the plot. The bounds of the BCI are denoted with vertical dashed lines. Each gene is plotted as a point. Lighter points with transparency are not significantly differentially expressed while solid red points denote significantly differentially expressed genes. Significant genes are labeled with only a random subset of genes labeled in the case of a wide BCI with too many to label. Q1 is excluded as the BCI spans the centromere and is very wide.

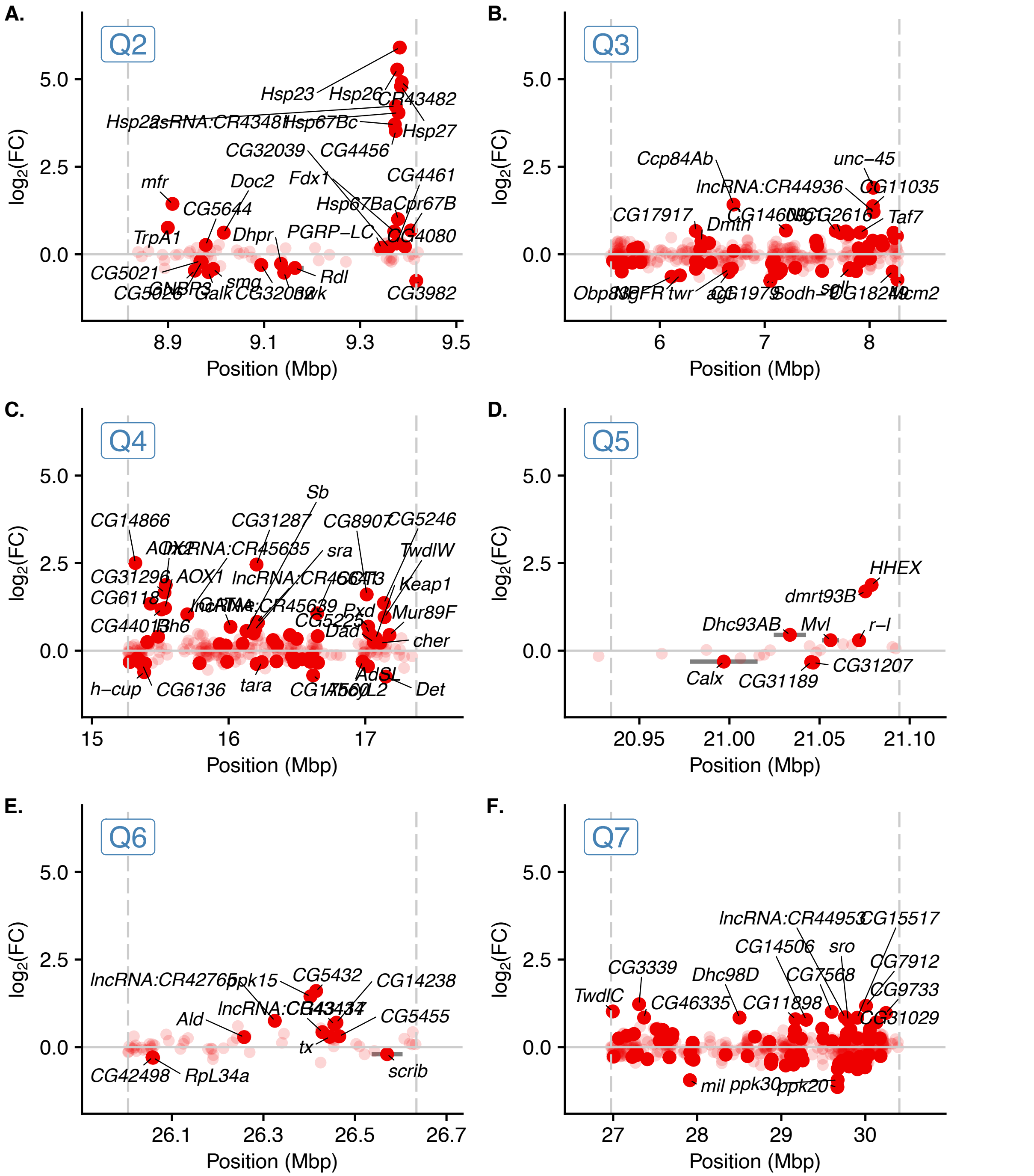

Figure S9: The $log_2$(fold change) between expression levels in incubated versus room temperature pools for each gene within each QTL BCI region. Each QTL id is labeled on the top left of the plot. The bounds of the BCI are denoted with vertical dashed lines. Each gene is plotted as a point. Lighter points with transparency are not significantly differentially expressed while solid red points denote significantly differentially expressed genes. Significant genes are labeled with only a random subset of genes labeled in the case of a wide BCI with too many to label. Q1 is excluded as the BCI spans the centromere and is very wide.

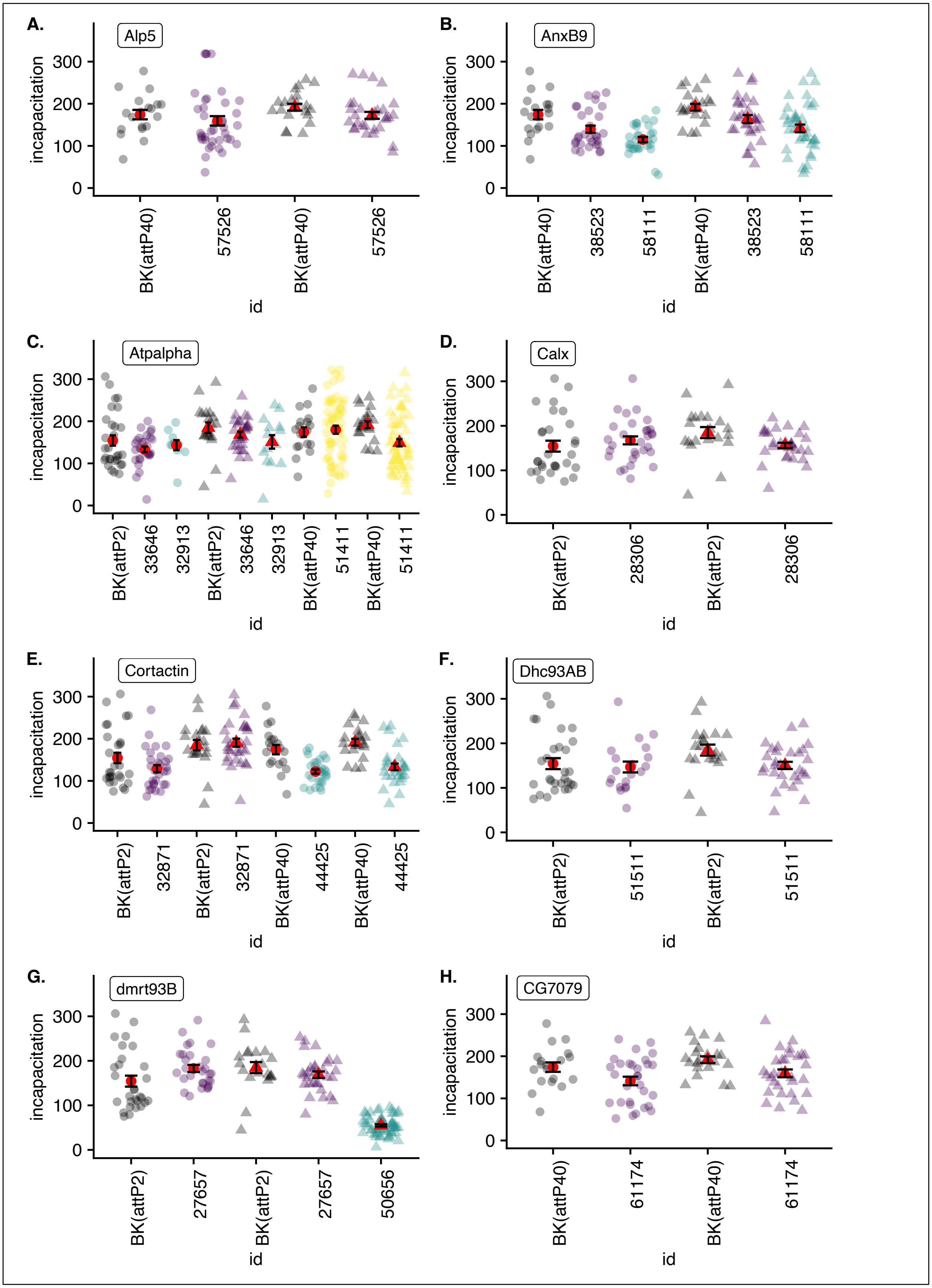

A.
Alp5
incapacitation
300
200
100
0
BK(attP40)
57526
BK(attP40)
57526
id
B.
AnxB9
incapacitation
BK(attP40)
38523
58111
BK(attP40)
38523
58111
id
C.
Atpalpha
incapacitation
BK(attP2)
33646
32913
BK(attP2)
33646
32913
BK(attP40)
51411
BK(attP40)
51411
id
D.
Calx
incapacitation
BK(attP2)
28306
BK(attP2)
28306
id
E.
Cortactin
incapacitation
BK(attP2)
32871
BK(attP2)
32871
BK(attP40)
44425
BK(attP40)
44425
id
F.
Dhc93AB
incapacitation
BK(attP2)
51511
BK(attP2)
51511
id
G.
dmrt93B
incapacitation
BK(attP2)
27657
BK(attP2)
27657
50656
id
H.
CG7079
incapacitation
BK(attP40)
61174
BK(attP40)
61174
id

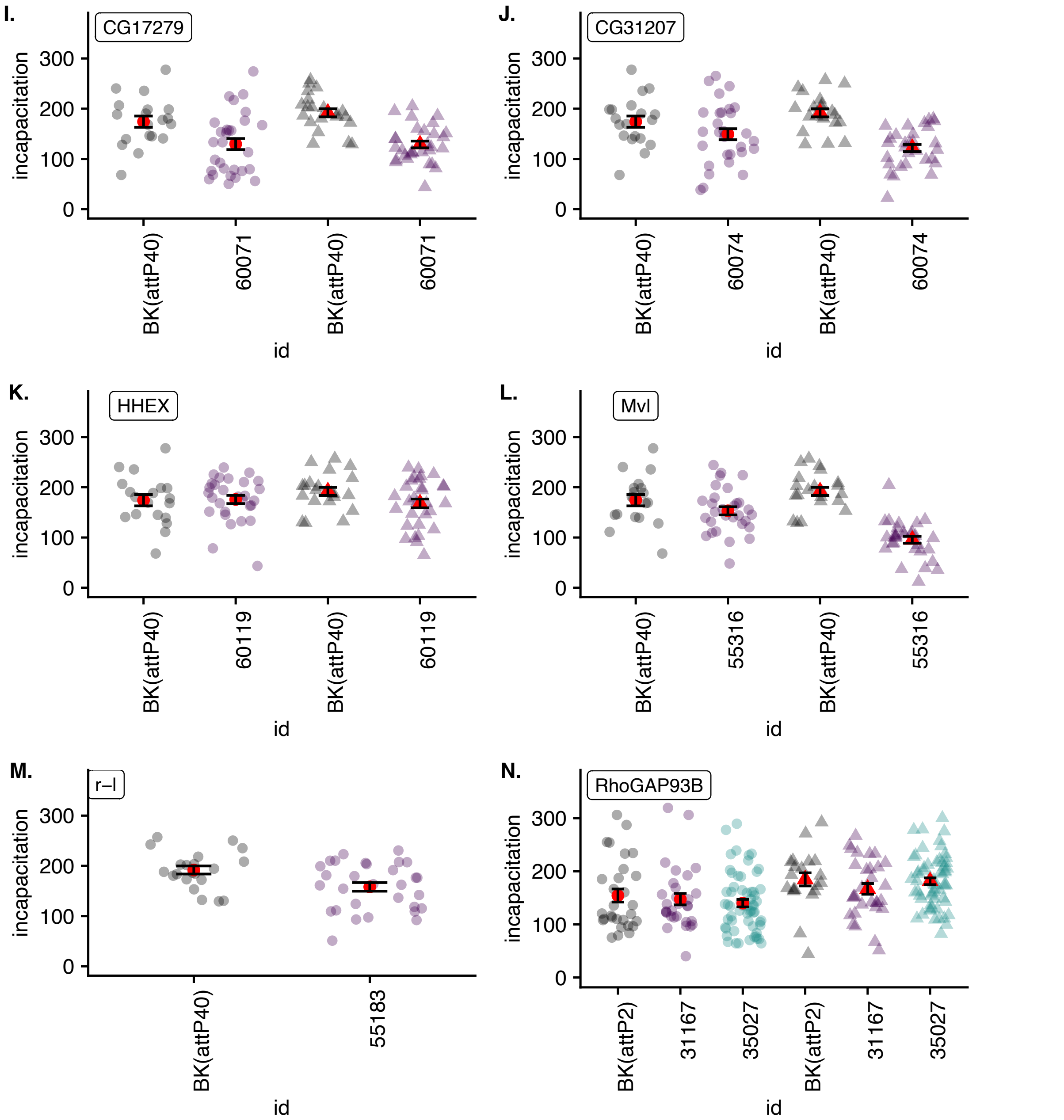


Figure S10: Incapcitation scores for each RNAi cross and background cross for each focal gene. Each point represents an individual fly. Background crosses for each RNAi type (attP2 or attP40) and driver (all cells vs all neurons) are shown in black and are to the left of each set of relevant RNAi crosses. Different drivers are shown with different shapes (all cells = circles; all neurons = triangles). Some focal genes are associated with more than one RNAi line with different lines denoted with different colors. The mean for each cross is shown in red with +/- 1 standard error overlayed in black.

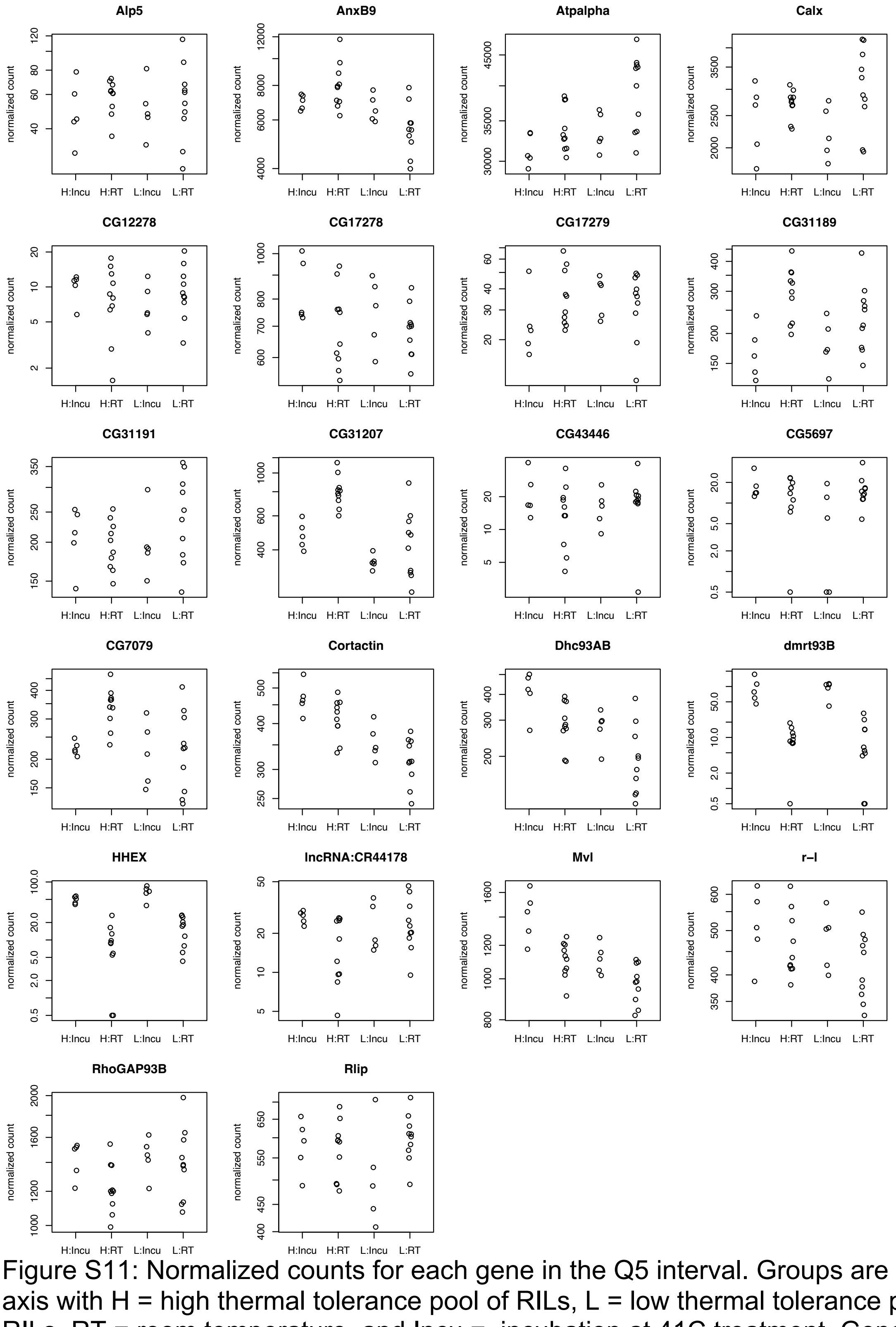


Figure S11: Normalized counts for each gene in the Q5 interval. Groups are on the x-axis with H = high thermal tolerance pool of RILs, L = low thermal tolerance pool of RILs, RT = room temperature, and Incu = incubation at 41C treatment. Gene labels are at the top of each plot.
